# NovumRNA: accurate prediction of non-canonical tumor antigens from RNA sequencing data

**DOI:** 10.1101/2024.11.05.622043

**Authors:** Markus Ausserhofer, Dietmar Rieder, Manuel Facciolla, Giorgia Lamberti, Rebecca Lisandrelli, Serena Pellegatta, Zlatko Trajanoski, Francesca Finotello

## Abstract

Non-canonical tumor-specific antigens (ncTSAs) can expand the pool of targets for cancer immunotherapy, but require robust and comprehensive computational pipelines for their prediction. Here, we present NovumRNA, a fully-automated Nextflow pipeline for predicting different classes of ncTSAs from patients’ RNA sequencing data. We extensively validated NovumRNA using publicly-available and newly-generated datasets, demonstrating the robustness of its analytical modules and predictions. NovumRNA analysis of colorectal cancer organoid data revealed comparable ncTSA potential for microsatellite stable and unstable tumors and candidate therapeutic targets for patients with low tumor mutational burden. Finally, our investigation of glioblastoma cell lines demonstrated increased ncTSAs burden upon indisulam treatment, and detection by NovumRNA of therapy-induced ncTSAs, which we could validate experimentally. These findings underscore the potential of NovumRNA for identifying synergistic drugs and novel therapeutic targets for immunotherapy, which could ultimately extend its benefit to a broader patient population.

## Introduction

Immunotherapy has revolutionized cancer treatment, shifting the focus from targeting malignant cells to supporting the patient’s own immune system in eliminating cancer cells^1^. Immunotherapy has achieved long-term durable responses in advanced cancers and is now approved worldwide for different tumor indications. Cancer immunotherapy broadly comprises immune checkpoint inhibitors (ICBs), personalized anticancer vaccines, and adoptive T-cell therapies^1^. Although different in their strategies, these modalities share the common goal of increasing the number and activity of T cells targeting tumor-specific antigens (TSAs)^2,3^. TSAs are mutated peptides that are presented to the immune system by major histocompatibility complex (MHC) molecules (called human leukocyte antigen, HLA, in humans) on the surface of tumor cells^3–5^. As they arise from protein aberrations present only in tumor cells and not in normal cells, TSAs can elicit strong immune responses mediated by T cells, which recognize these peptides as “non-self”. The most well-characterized class of TSAs are *neoantigens*, namely mutated peptides derived from non-synonymous single nucleotide variants (SNVs) and insertions or deletions (indels). TSAs are considered the major targets of immunotherapy responses and their computational prediction in cancer patients is the basis for personalized adoptive T-cell therapy and anticancer vaccination^3,6,7^. Therefore, the computational identification of personalized TSAs from patient’s next-generation sequencing (NGS) data is a key task in immune-oncology.

Despite the success and potential of ICBs, the majority of patients still do not benefit from immunotherapy: either they are not eligible for it or do not respond^8,9^. Therefore, there is a pressing need to identify a larger pool of TSA targets to extend the population of cancer patients treated with immunotherapy, and to understand and reverse the mechanisms of resistance to augment its clinical success. Tumor mutational burden (TMB), defined as the number of total non-synonymous mutations in a tumor, is a major predictive biomarker for immunotherapy response as TMB-high tumors are more likely to generate neoantigens^7,10–13^. Nevertheless, the correlation between TMB and immunotherapy outcome is far from perfect, and patients with low TMB have also shown responses to immunotherapy^14^. For instance, renal cell cancer, despite having low TMB, has shown a good response rate to ICBs (∼25%)^14^. Neoadjuvant ICBs therapy has led to comparable pathological responses in mismatch-repair-deficient and -proficient early-stage colorectal cancers (CRC)^14,15^.

Altogether, these results suggest that immunotherapy can elicit immune responses targeting TSAs generated from additional aberrations that do not fall into the canonical definition of TMB. Recent studies have shown that TSAs can also arise from the expression of cancer-specific alternative splicing events^16^, gene fusions^17^, retroviral elements^18^, and non-coding regions^19,20^. These non-canonical TSAs (ncTSAs) can represent an additional pool of therapeutic targets whose potential is yet to be fully uncovered^19–22^. Therefore, ncTSAs hold the promise to extend the benefit of immunotherapy to a wider population of cancer patients, possibly targeting also low-TMB tumors, but their prediction from tumor NGS data requires ad hoc computational pipelines.

While there are several tools available for the prediction of neoantigens from SNV and indels, non-canonical TSA prediction is still in its early stages. To date, there is no consensus on best practice and workflows for ncTSA prediction, and third-party benchmarking studies are lacking. However, computational pipelines and workflows have been recently proposed to predict certain classes of ncTSAs from patients’ NGS data^19,23–32^. These methodologies founded a new toolkit for ncTSA-prediction, with each approach offering specific strengths. However, none of these methods satisfies altogether a set of criteria that are required to warrant their broad adoption in research and clinical settings. First, providing comprehensive metadata about the predicted ncTSAs regarding their origin, genomic annotation, and, most importantly, likelihood to be presented to and recognized by T cells –information which is crucial for prioritization and experimental validation. Second, predicting ncTSAs from different types of non-canonical sources, as the best targets can depend on the specific tumor. Third, including predictions not only for class-I but also class-II ncTSAs, which are the targets of CD4^+^ T cell-mediated anticancer immunity^33,34^. Fourth, implementing a dynamic and dedicated filtering approach to minimize false positives, namely *self-peptides* that are also expressed in non-cancerous tissues. Finally, offering an efficient, standalone, and well-documented software solution that can handle raw data inputs (without relying on third-party analytical tools), while ensuring ease of use and installation.

Here, we present NovumRNA: a Nextflow^35^ pipeline for the prediction of ncTSAs from raw tumor RNA sequencing (RNA-seq) data, addressing all the desired features outlined above. We validated NovumRNA through the analysis of different RNA-seq datasets, demonstrating the robustness of its different analytical modules and final predictions. We further applied it to the analysis of a new RNA-seq dataset generated from ten tumor organoids derived from patients with microsatellite stable (MSS) and microsatellite instable (MSI) CRC. With our analysis, we show how large control datasets can be used to more stringently ensure the tumor-specificity of the predicted ncTSA. Moreover, this analysis revealed a comparable ncTSA burden for both MSS and MSI patients, and the identification of shared candidates across multiple patient-derived organoids, underlying the importance to further investigating the landscape of ncTSAs in hard-to-treat cancers. Finally, we generated an RNA-seq dataset from four patient-derived glioblastoma cell lines treated with the splicing-modulating drug indisulam. Using NovumRNA to analyze the data, we found a profound remodeling of the tumor-cell transcriptomes induced by this drug, which resulted in the generation of ncTSAs, induced by the treatment. Using co-culture experiments, we experimentally validated NovumRNAs predictions, demonstrating its accuracy as well as its value for potentially pinpointing synergistic partners for immunotherapy.

## Results

### The NovumRNA pipeline allows the prediction of non-canonical tumor antigens from RNA sequencing data

NovumRNA is a multi-step, fully-automated pipeline for the prediction of ncTSAs from human tumor RNA-seq data (**Fig. 1**). It takes single- or paired-end RNA-seq reads (FASTQ files) as input, which are then aligned to the reference genome using either HISAT2^36^ or, alternatively, STAR^37^, followed by reference-guided transcript assembly with StringTie^38^.

**Figure 1.**
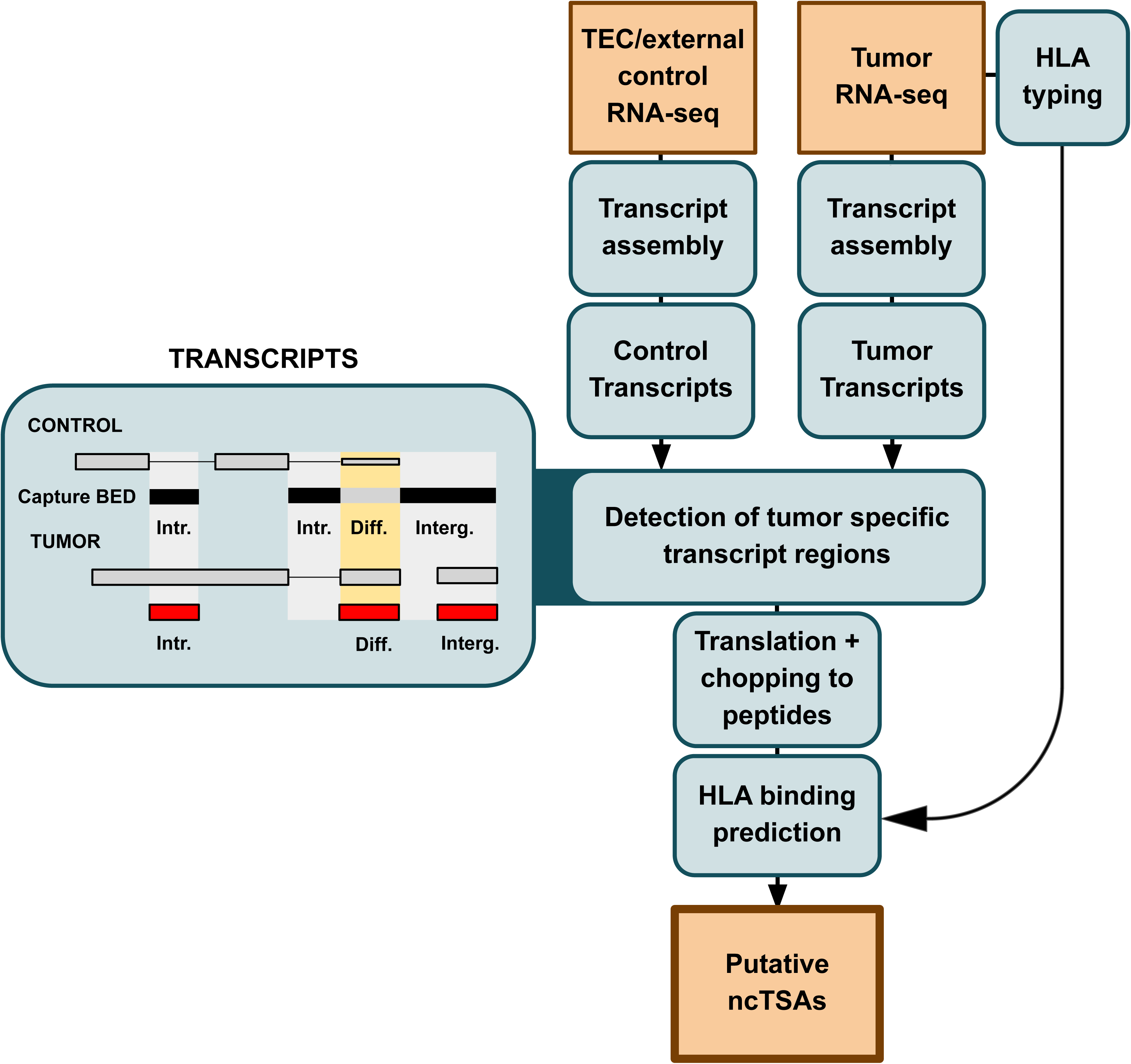
Schematization of the NovumRNA framework. Tumor RNA sequencing FASTQ files (and, in parallel, non-cancerous control samples) are aligned to the reference genome to perform reference-guided transcript assembly. Tumor-specific and differentially expressed transcript fragments are identified by comparison with the control transcripts stored in the form of a capture BED text file (derived from the internal thymic epithelial cell (TEC) reference or from external, user-provided control data). Filtered, tumor-specific transcript fragments are translated and fragmented into peptides of specified lengths. In parallel, patient-specific class-I and -II human leukocyte antigen (HLA) molecules are identified. Tumor-specific peptides are tested for their binding affinity towards the patient’s HLA class I and class II molecules. A final output data frame is generated containing all relevant metadata from previous steps, including additional endogenous retrovirus (ERV) annotation, to select the best non-canonical tumor specific antigen (ncTSA) candidates.

NovumRNA seeks to identify *tumor-specific* transcript fragments, defined as genomic regions that are exclusively covered by transcripts identified from the input tumor RNA-seq data and are instead completely absent in transcripts assembled from control data obtained from non-cancerous tissues. These control regions are identified by NovumRNA and stored in a capture BED file (details in Methods). By default, the regions annotated in the capture BED file are derived from an internal control RNA-seq database, which is based on 32 RNA-seq samples from human thymic epithelial cells (TECs). As demonstrated in a previous study^19^, TECs can be used as “normal control” for the selection of TSAs thanks to their unique capability to express most known genes and for their unique role in inducing T-cell central tolerance to self-peptides. Alternatively, users can build their own filtering database, providing NovumRNA with a selection of control RNA-seq samples. Strictly tumor-specific transcript fragments are considered as *novel* and annotated by their genomic location as *intronic* or *intergenic*. Fragments which have high expression levels in tumor cells but have low expression levels in healthy controls are also retained for prediction and annotated as *differential*.

Transcript fragments that pass the filtering steps (with user-customizable cut-offs) are then translated into peptide sequences. NovumRNA translates the full transcript of origin, either using the corresponding reference protein based on GENCODE annotation^39^, when available, or by selecting the reading frame based on the presence/location of methionine and/or Kozak motifs (details in Methods). The translated fragments are then chopped into peptides of a specified length, which are additionally filtered against the GENCODE reference proteome to make sure they are not present within non-tumor-specific proteins.

In parallel, the patient’s class-I and -II human leukocyte antigen (HLA) molecules are identified from RNA-seq data using OptiType^40^ and HLA-HD^41^, respectively. Alternatively, NovumRNA allows specifying the patient’s HLA types to be used for ncTSA prediction (skipping HLA typing). Finally, tumor-specific peptides are tested for their binding affinity towards the patient’s class-I and -II HLA molecules using netMHCpan and netMHCIIpan^42^, respectively.

NovumRNA reports ncTSAs from intronic, intergenic, and differential origin. These ncTSAs are further categorized as endogenous retrovirus (ERV)-derived based on their overlap with known ERV regions according to HERVd annotation^43^.

In addition to HLA-peptide binding affinity, NovumRNA provides a rich set of ncTSA-specific features relevant for prioritization and validation, including: exact genomic location of the predicted peptide, expression level of the original transcript, and different read-coverage statistics. Additionally, NovumRNA calculates a novel metric, called “TPM_iso_perc”, akin to the concept of clonality. It quantifies the fraction of TPM expression explained by the transcript from which a ncTSA derives from, relative to the sum of TPM expression across all other transcript isoforms. A high “TPM_iso_perc” value indicates a higher “clonality” of the underlying non-canonical tumor aberration, a feature that is key to ensuring the selection of high-quality ncTSAs having a higher immunogenicity potential^2,44^.

NovumRNA is implemented in Nextflow DSL2^35^ and uses Singularity containers^45^ to allow for easy installation, parallelization, and customization. For additional information on NovumRNA methodology and usage, we refer to the Methods section and to its GitHub repository (https://github.com/ComputationalBiomedicineGroup/NovumRNA).

### Comprehensive analysis of public datasets demonstrates the robustness of NovumRNA predictions

To ensure the accuracy of the RNA-seq-based HLA typing module implemented in NovumRNA, we performed a benchmarking analysis of four state-of-the-art tools: arcasHLA^46^, HISAT^47^, HLA-HD^41^, and OptiType (only class I)^40^. Building upon our previous work^24^, we constructed a gold standard by combining data from two studies from the 1000 Genomes Project^48–50^. In our assessment, all tools displayed high accuracy for both class-I and -II genes, confirming that HLA typing solely based on RNA-seq data is reliable (**Fig. 2A**). HLA-HD and OptiType obtained the best performance on all class-I HLA genes, with percentage of correct calls ranging in 97.5-99.6 for HLA-HD and 98.2-99.2 for Optitype. Notably, Optitype never resulted in a missing call (while HLA-HD has 0.4% missing calls); this feature has paramount importance for personalized cancer immunotherapy, where putative ncTSAs need to be predicted for every patient. For this reason and for its open-source licensing scheme, we based the class-I HLA typing module of NovumRNA on OptiType. On class-II HLA genes, HLA-HD performed best, with correct calls ranging between 99.2-99.6 (0.4% missing calls); therefore class-II HLA typing in NovumRNA is performed with HLA-HD.

**Figure 2.**
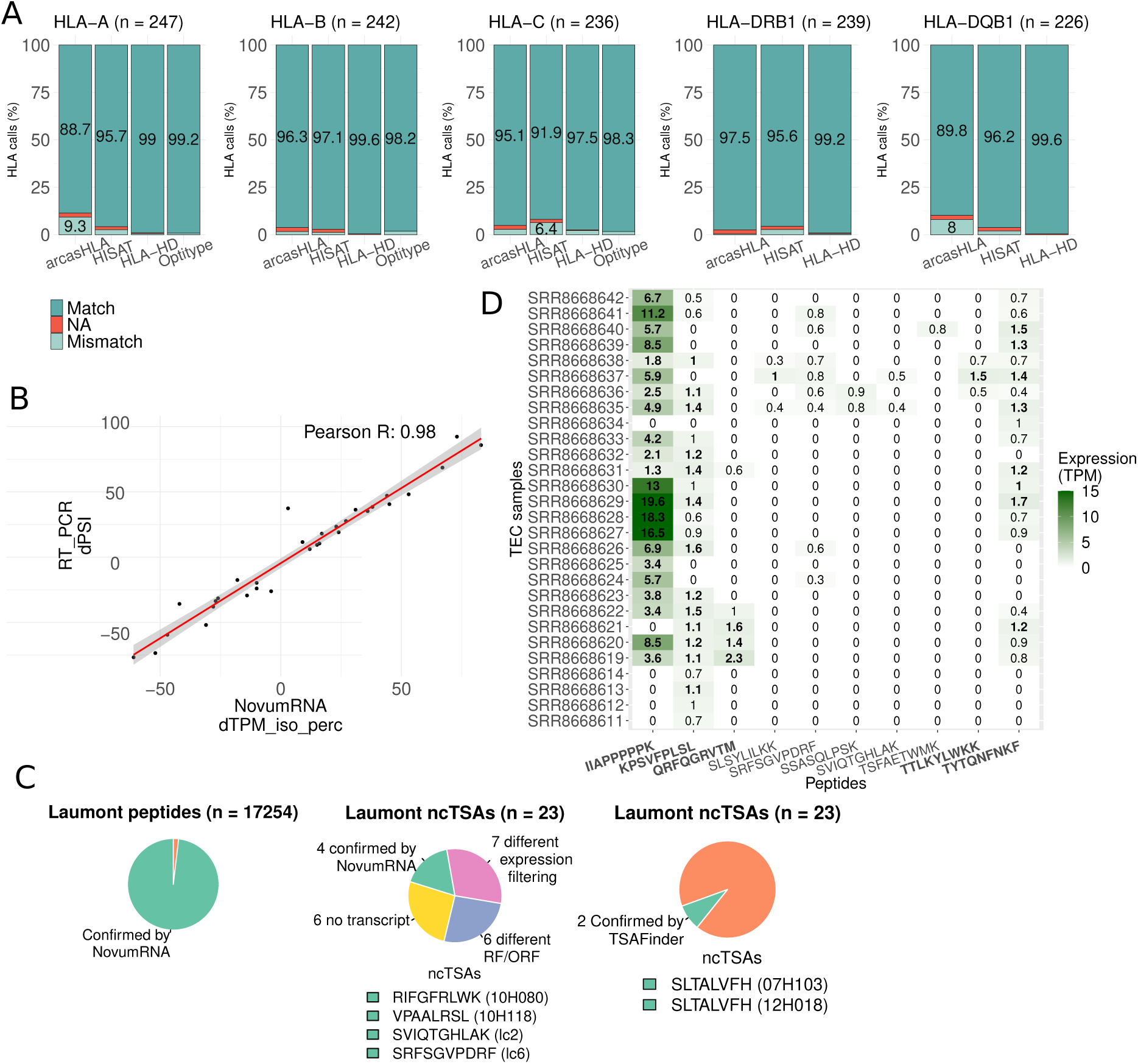
NovumRNA validation. **A)** Percentage of correct (Match), incorrect (Mismatch), and missing (NA) HLA calls inferred by HLA-HD, HISATgenotype, arcasHLA (class-I and II) and Optitype (for class-I genes), evaluated on RNA-seq data from the 1000 Genomes project. **B)** Validation of NovumRNA “TPM_iso_perc” estimates using RNA-seq data from six cancer cell lines from Shen et al. ^52^. Pearson correlation of delta exon inclusion rates from 32 out of 34 validated cassette-exons estimated by NovumRNA (“TPM_iso_perc”, x-axis) vs. reverse transcription polymerase chain reaction (RT-PCR)-determined values (“y-axis”). **C)** Validation of peptide and non-canonical tumor specific antigen (ncTSA) predicted with NovumRNA from Laumont et al. ^19^. RNA-seq data. Left-to-right: pie charts showing the fraction of peptides confirmed by NovumRNA (in green, 96.6 %); number of ncTSAs confirmed (in green, with reported peptide sequence and sample identifier) and not confirmed by NovumRNA, or by TSAfinder. **D)** Heatmap showing the expression in transcripts per millions (TPM) in the control thymic epithelial cells (TEC) samples of peptides identified as TSA in the Laumont study. A “/” in the peptide sequence represents alternative amino acids (A/B), since the underlying nucleotide sequence showed a single nucleotide polymorphism (SNP), or mutation at this position.

To assess NovumRNAs effectiveness in detecting and quantifying differentially-spliced events from RNA-seq data –a daunting task even for specialized splicing-detection tools^51^– we considered a publicly available RNA-seq dataset from pancreatic cell lines, for which 34 cassette-exon events were experimentally confirmed via reverse transcription polymerase chain reaction (RT-PCR)^52^. NovumRNA confirmed 32 out of the 34 events, and its “TPM_iso_perc” estimates showed a high correlation with the differential percent spliced inclusion (dPSI) values based on RT-PCR measurements performed in the original study (Pearson correlation of 0.98, p = 3×10^−21^) (**Fig. 2B**).

To assess NovumRNAs performance in ncTSA prediction, we used data from a study where putative ncTSAs were predicted from seven primary human tumor samples using a proteogenomic approach based on matched RNA-seq and mass spectrometry (MS) data^19^. In this study, Laumont and colleagues made available for each sample, besides the transcriptomic data, the whole list of MS eluted peptides (n = 17,254), including ncTSAs (n = 23) identified via integrative analysis of RNA-seq and MS data. We applied NovumRNA to the tumor RNA-seq data, considering peptide lengths of 8-11 amino acids and using default settings and references.

We first compared class-I HLA types predicted with OptiType by NovumRNA with those reported in the Laumont study (**Table 1**). All HLA types agreed at four-digit resolution, except for one HLA allele for sample 10H118, which agreed on only at two-digit resolution (HLA-C*07:18/07:37 NovumRNA vs. HLA-C*07:01 Laumont) (**Table 1**).

**Table 1.** Eluted peptides from Laumont et al. confirmed by NovumRNA. Shown are the total number of mass spectrometry (MS)-confirmed, eluted peptides from seven primary human cancer samples, as well as the number and percentage of peptides confirmed by NovumRNA, together with the proportion of matching HLA type calls.

| Sample | HLA match | Peptides Laumont | Confirmed by NovumRNA |
| --- | --- | --- | --- |
| 07H103 | 5/5 | 804 | 776 (96.5%) |
| 10H080 | 4/4 | 3111 | 3045 (97.9%) |
| 10H118 | 3/4 | 171 | 155 (90.6%) |
| 12H018 | 6/6 | 372 | 357 (96.0%) |
| lc2 | 5/5 | 5289 | 5202 (98.4%) |
| lc4 | 4/4 | 4617 | 4551 (98.6%) |
| lc6 | 4/4 | 2890 | 2838 (98.2%) |

We further compared NovumRNA-predicted peptides to the list of MS-confirmed, eluted peptides. Since NovumRNA, by default, only translates transcripts containing relevant tumor-specific fragments, the translation module was run separately from the pipeline on all transcripts assembled by StringTie, and the resulting proteins were fragmented into all possible 8-11 amino acid-long peptides. On average, 96.6% of eluted peptides were confirmed by NovumRNA for each sample (**Table 1** and **Fig. 2C** left).

Finally, we compared the putative ncTSAs identified by NovumRNA (predicted solely from RNA-seq data) with the ones reported in the original study (using matched RNA-seq and MS data). From the total 23 ncTSAs identified in the Laumont study, 4 were confirmed also by NovumRNA, including confirmation of the correct HLA type (**Table 2** and **Fig. 2C** middle). For comparison, a recent study focused on the prediction of ncTSAs from this dataset using a tool called TSAFinder^25^ could identify only a single matching peptide, present in two samples (SLTALVFHV, samples 07H103 and 12H018) (**Fig. 2C** right).

**Table 2.**
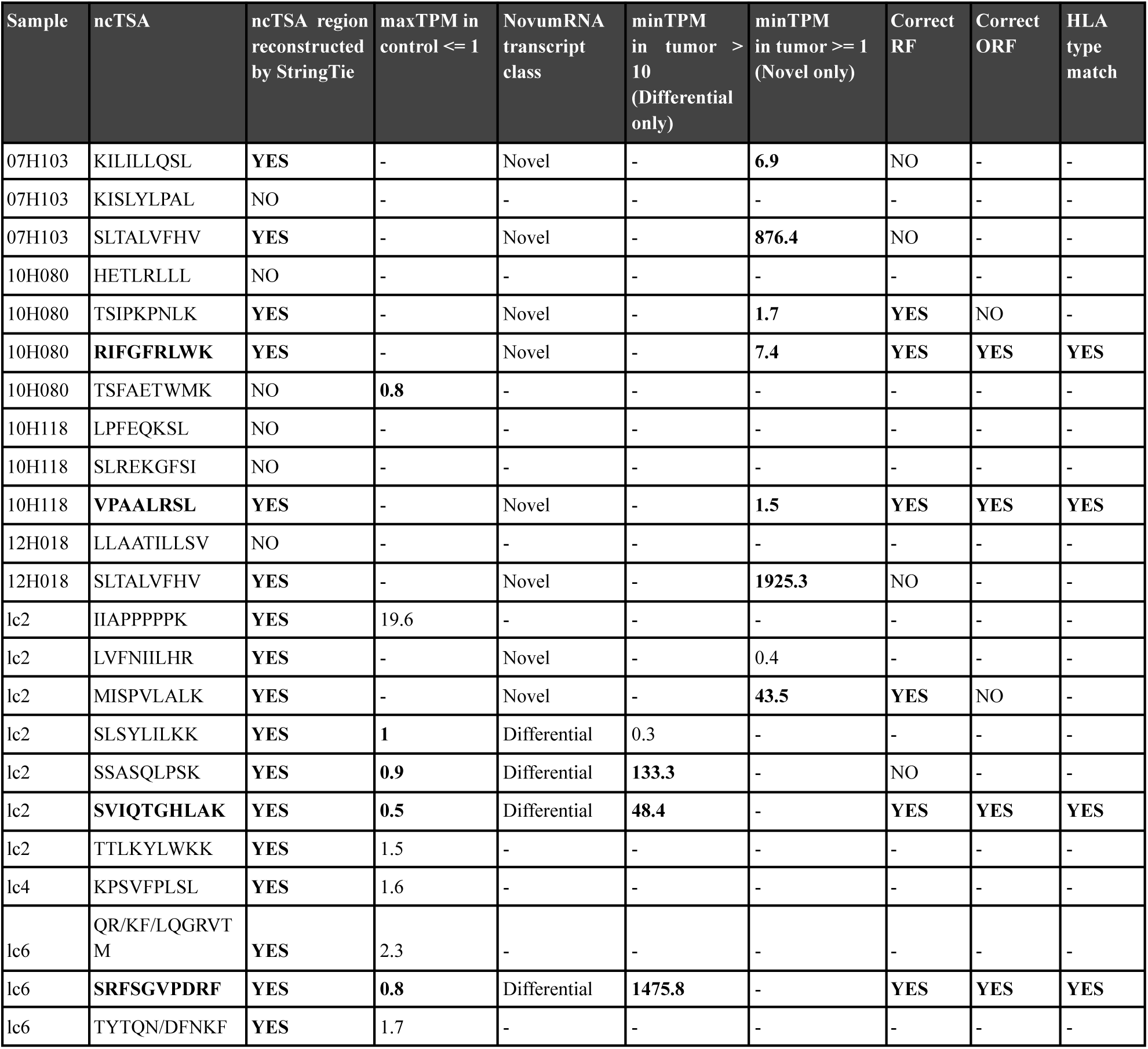
Comparison of NovumRNA and Laumont ncTSA predictions. Table shows the features of the ncTSAs identified in the Laumont study, which were confirmed (in bold) or not confirmed by NovumRNA according to the adopted filtering approach. The “ncTSA region”column indicates whether the ncTSA region was reconstructed by StringTiet. The “maxTPM” column indicates wether the maximum expression in the control samples did not exceed 1 TPM.The “NovumRNA class” reports NovumRNA-determine ncTSAclasses, as “ovel” or “ifferential”. For “differential” ncTSAs, the minimum level of expression in (“minTPM”) in the tumor sample must be greater than 10TPM. For “novel” ncTSAs, “minTPM” should be greater than or equal to 1 TPM. The “Correct RF” and “Correct ORF” columns indicats if the correct reading frame (RF) and open reading frame (ORF) was predicted by NovumRNA. Finally, the “HLA type match” column indicates if the peptide-bound human leukocyte antigen (HLA) type predicted by NovumRNA matches that identified in the study A. A “/” in the peptide sequence represents an alternative amino acid, since the underlying nucleotide sequence showed a point mutation at this position.

Given the differences in terms of used input data and analytical approach, we investigated at which point of the analysis NovumRNA and Laumont ncTSA predictions diverged. As the location (chromosome plus start and stop position) of all the 23 ncTSAs were available, we checked if these genomic regions were covered by the transcripts assembled by the StringTie module of NovumRNA (**Table 2**). Of the 23 ncTSA regions, 17 were successfully reconstructed, while six were undetected (**Table 2** and **Fig. 2C** middle).

For an ncTSA to be classified as “tumor-specific” by NovumRNA, its maximum expression across control samples must not exceed a certain threshold, set at 1 transcript per million (TPM) by default. Despite being labeled as “tumor-specific” in the original study, we found seven ncTSAs to be expressed in the TEC controls (**Fig. 2D**). Three of them showed non-negligible expression in multiple samples, exceeding the 1 TPM cut-off: IIAPPPPPK (max: 19.6 TPM), KPSVFPLSL (max: 1.6 TPM) and TTLKYLWKK (max: 1.5 TPM) (**Table 2**, **Fig. 2D**). Given these expression levels in the controls, NovumRNA did not select these three peptides as ncTSAs (**Table 2**). Two more ncTSAs, QR/KF/LQGRVTM (max: 2.3 TPM) and TYTQN/DFNKF (max: 1.7 TPM), this time with a reported expression in the controls also in the original study, also exceeded NovumRNA cut-offs and were filtered out. From the remaining peptides, one differential (SLSYLILKK) and one novel (LVFNIILHR) putative ncTSA were instead discarded by NovumRNA due to their low expression in the tumor samples (**Table 2**).

Finally, six of the remaining putative ncTSAs were not identified by NovumRNA due to a different prediction of their reading frame (RF) or open reading frame (ORF). Of note, NovumRNA was able to predict the correct RF for 6 over 10 peptides, and the correct ORF for 4 over 10 peptides (**Table 2**), without relying on 3-frame translation or additional proteomics data as done in the original study. Due to the complexity of RF and ORF identification from de novo-assembled transcripts, we tested an alternative approach based on BORF, a tool specifically designed for ORF prediction^53^. The usage of BORF in the NovumRNA framework led to the identification of one additional ncTSA (SLTALVFH, present in two samples), counterbalanced by the loss of one ncTSA found by NovumRNA (RIFGFRLWK).

Taken together, the results of our benchmarking efforts based on several publicly-available datasets underscore the robustness of NovumRNA analytical modules and overall approach.

### Prediction of non-canonical tumor antigens in colorectal cancer organoids identifies potential targets in microsatellite-stable tumors

NovumRNAs filtering approach leverages a small but effective internal database of TEC-derived control transcripts to discriminate and filter out transcript sequences that are not tumor-specific. Depending on the specific application, larger and/or tissue-specific control data might be preferred. NovumRNA offers a mode called “capture_bed”, where user-supplied RNA-seq data from healthy tissues can be used to create a new capture BED file, substituting the default TEC control database.

To demonstrate the value of this approach, we analyzed an RNA-seq dataset generated from ten CRC organoids (**Table S1**) using the default TEC filtering strategy, as well as an approach based on a capture BED file built from healthy colon RNA-seq data from the Genotype-Tissue Expression (GTEx) project^54^. To assess the impact of the GTEx-based filtering approach, we performed a saturation analysis by randomly selecting an increasing number of GTEx colon samples for building the control database. In brief, the analysis was run with capture BED files made from 10-to-260 GTEx colon samples randomly-selected from the whole pool of 262 (**Table S2**), repeating each sampling and corresponding analysis 10 times to also assess the robustness of the predictions (Methods).

As expected, the average number of predicted ncTSAs decreased as the number of healthy colon samples used for filtering (referred in the following to as “filter-sample size”) increased, with similar patterns across different organoids (**Fig. 3A**). The default NovumRNA approach (based on 32 TEC samples) predicted on average 70% more peptides compared to the GTEx solution at 30 samples (**Fig. 3A**). The coefficient of variation (CV), indicating the variability of the predictions across the ten replicates, was moderate when only ten GTEx samples were used (average 27%), but dropped rapidly already at 20 samples (average 8%) and decreased further as the filter-sample size grew **(Fig. 3B**).

**Figure 3.**
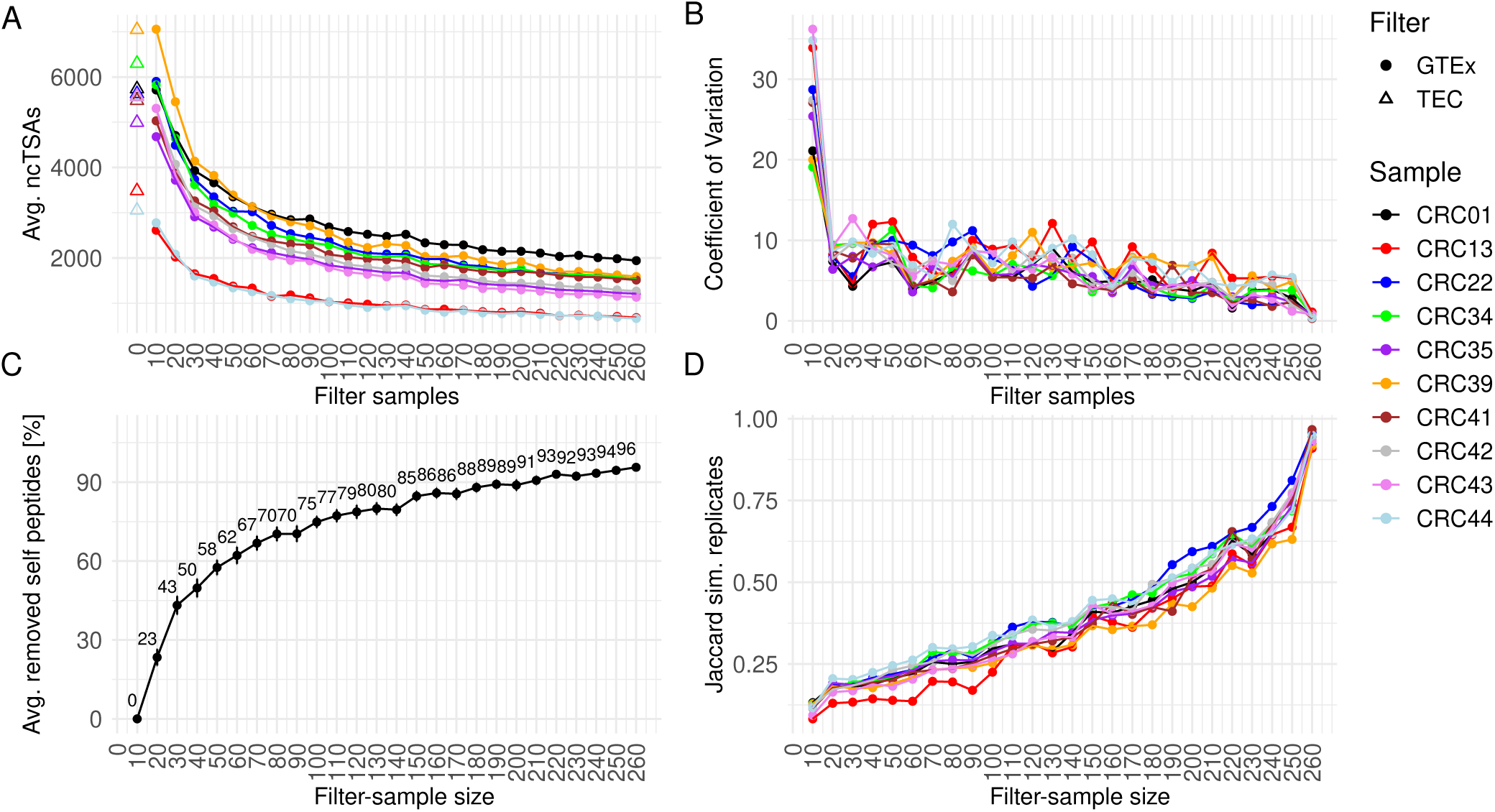
Effect of control database size on NovumRNA predictions on RNA sequencing data from colorectal cancer organoids. In all panels, the x-axis shows number of healthy colon GTEx samples used for filtering **A)** Average number of predicted non-canonical tumor-specific antigens (ncTSAs). The triangles indicate the solution obtained using the default TEC reference for filtering. **B)** Coefficient of variation across replicates for the statistics shown in panel A. **C)** Percentage of filtered self peptides, considering the 10-sample filtering step as reference (i.e. 0% removal rate by definition). Dots represents the average across all organoids and replicates, while the vertical bars indicate the standard deviation. **D)** Jaccard similarity of the predicted ncTSAs across all ten replicates for each organoid and filtering step.

To evaluate the benefit of increasing the filter-sample size, we calculated the percentage of removed self-peptides at every filtering increment. Briefly, for every prediction, we identified putative self-peptides as the predicted ncTSAs which were not confirmed by the results obtained using the full GTEx dataset for filtering. Considering the 10-sample filtering step as a reference, we calculated the percentage of removed self-peptides at every filter-sample size, averaging across replicates and organoids (details in Methods). Overall, a filter based on just 30 GTEx samples (i.e. ∼12% of the total GTEx data) already resulted in the removal of 43% of self peptides, while at 150 samples (∼58%), a removal rate of 85% was achieved. (**Fig. 3C**). To further assess filtering dynamics, we counted, for each filtering step, the number of ncTSAs unanimously predicted across all replicates, and quantified the robustness of the solution using the Jaccard similarity score. Starting with an average value of 0.11 across all organoids, similarity increased until 0.93 at the maximum filtering-sample size (**Fig. 3D**).

We then considered NovumRNA predictions obtained using all 262 GTEx controls to gain an overview of ncTSA landscape in the CRC organoids. Overall, there were no significant differences between MSI and MSS tumors in the total number of novel ncTSAs (p = 0.904, **Fig. 4A**). Similarly, there were no significant differences for the individual ncTSA classes (**Fig. S1**). The top ncTSA candidates, selected based on HLA binding affinity, transcript expression and coverage, and clonality likelihood (details in Methods), represented all novel classes (**Fig. 4B** and **Table S3**). By querying the identified peptides in public antigen databases, we found matches for 11 novel and 19 differential ncTSAs in four databases^55–57^ (**Fig. 4B** and **Table S4**). We could further assign the sequences of six ERV-derived ncTSAs (TQSLFGGLF, SLFGGLFTR, LCSMRKIHL, ENYSQRGLF, CSMRKIHLR, FRLFLIRPH), three of which were shared across two organoids, to the ERVH-5 retroviral gene.

**Figure 4.**
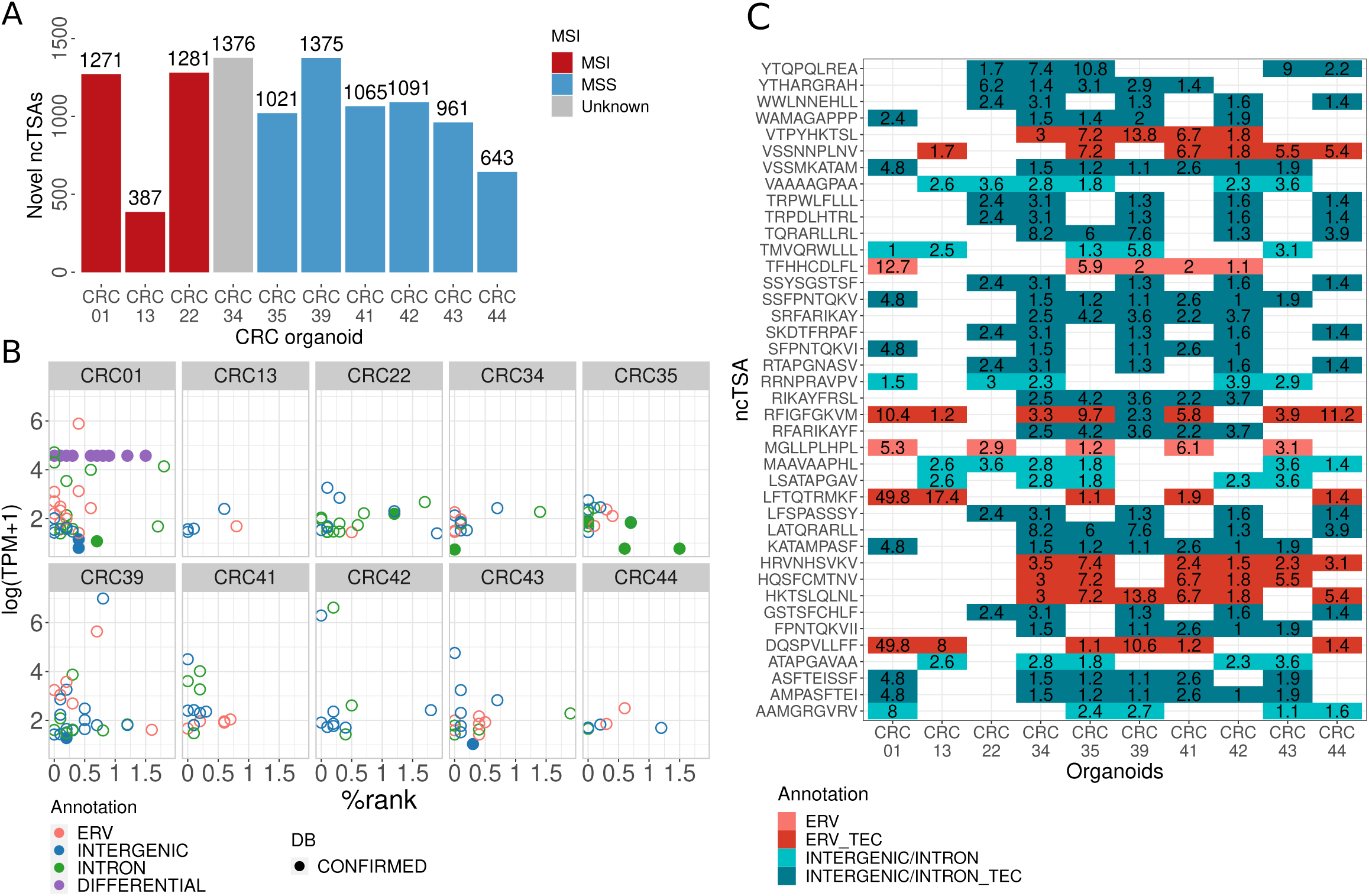
NovumRNA analysis of colorectal cancer organoids. **A)** Number of novel non-canonical tumor-specific antigens (ncTSAs) in ncTSAs in microsatellite instable (MSI) and microsatellite stable (MSS) colorectal cancer (CRC) organoids. **B)** Top ncTSA candidates predicted by NovumRNA for all ten organoids, with reported binding affinity as % rank (x-axis), expression level as log-scaled transcripts per millions (TPM) (y-axis), ncTSA class (indicated by the dot color). Database hits are indicated as full dots. **C)** Heatmap showing the top-shared ncTSAs across CRC organoids (in at least five organoids). Tiles are colored by ncTSA origin annotation; darker colors indicate wether the ncTSA were also found using the default thymic epithelial cells (TEC) filtering approach. The numbers in tiles indicatethe TPM expression of the transcript the ncTSAs derive from.

We further investigated if the predicted ncTSAs were shared across CRC organoids. 834 ncTSAs were shared across at least two organoids, while only one was shared across eight organoids (RFIGFGKVM); no ncTSAs were shared across all ten organoids (**Fig. 4C** and **Table S5**). Notably, 159/834 shared ncTSAs (19%) were derived from ERV sequences, including the peptide shared across eight organoids. Among all ncTSAs predicted using the whole GTEx dataset for filtering, 8,161 (65%) were also present among the final ncTSAs predicted using the default TEC filtering approach.

### NovumRNA identifies non-canonical tumor neoantigens induced by indisulam treatment in glioblastoma cell lines

RNA-seq provides a cost-effective technique to investigate the dynamics of gene expression upon perturbation. NovumRNA can be effectively applied to perturbed RNA-seq data generated from treated cancer cell lines to investigate how drugs modulate the expression of ncTSAs, potentially influencing the immunogenicity of tumor cells. To test NovumRNA on this application, we analyzed an RNA-seq dataset generated from the human bladder cancer cell line UM-UC-3 treated with the methylation inhibitor 5-Aza-2′-deoxycytidine (5-Aza-CdR) and subjected to RNA-seq after 5, 9, 13, and 17 days of treatment^58^. NovumRNA was applied with default parameter settings, considering binding peptides of 9 amino acids in length (Methods). The highest number of novel ncTSAs was predicted at day 17 (**Fig. S2A**), in accordance with the largest drug-induced transcriptional effect seen in the original study^58^. To rule out the possibility that this pattern was due to sequencing depth bias, we computed Pearson correlation between the number of mapped reads and the total number of ncTSAs predicted per treatment, confirming no association (R = 0.25, p = 0.68, **Fig. S3**).

We analyzed a second dataset from cell lines treated with the splicing-modulating drugs indisulam and MS023^59^. In this study, RNA-seq data was generated from three human melanoma cell lines (510 MEL, A375, SK MEL 239), with triplicates for each cancer cell line, treated for 96 hours with indisulam, MS023, or with dimethyl sulfoxide (DMSO), used as a control.

NovumRNA analysis revealed an increase in the number of predicted novel ncTSAs (i.e., intronic, intergenic, and ERV) in all three cell lines treated with indisulam compared to the control group (**Fig. S2B**). The number of ncTSAs of differential origin showed instead no change (510 MEL) or a decrease (SK_MEL_239, A375) upon treatment. Compared to the A375 cell line, which showed moderate changes upon indisulam treatment, the SK_MEL_239 and, especially, 510 MEL cell lines exhibited a marked augmentation in the number of predicted ncTSA in the treated vs. control condition: SK_MEL_239: intronic (+150%, p = 0.03), intergenic (+140%, p = 0.05), ERV (+230%, p = 0.05); 510 MEL: intronic (+380%, p = 0.05), intergenic (+230%, p = 0.05), ERV (+330%, p = 0.05). Treatment with MS023 resulted in no significant changes in terms of ncTSA numbers, with the exception of intronic ncTSAs in the 510 MEL cell line (+118%, p = 0.05) (**Fig. S2B**).

The profound changes in ncTSA potential fostered by indisulam treatment motivated us to investigate the effect of this drug in a non-immunogenic cancer: glioblastoma. Primary cell lines were generated from surgical material collected from four distinct glioblastoma patients and grown in vitro as neurospheres (GB-NS). GB-NS were then treated with indisulam or vehicle control, and subjected to bulk RNA-seq (Methods). The resulting data was analyzed with NovumRNA, using 120 healthy brain GTEx samples as a normal control for filtering (**Table S2**).

Indisulam treatment of GB-NS cells resulted in an increase of predicted novel ncTSAs (ERV, intergenic, and intronic) across all four samples, but with peaks that varied in latency depending on the patient’s sample (**Fig. 5B**). Also the magnitude of change upon indisulam treatment was patient-specific: the changes in the number of predicted ncTSA were particularly marked for patient 1 and, limited, instead for patient 3. Differential ncTSAs decreased across all patients upon indisulam treatment.

**Figure 5.**
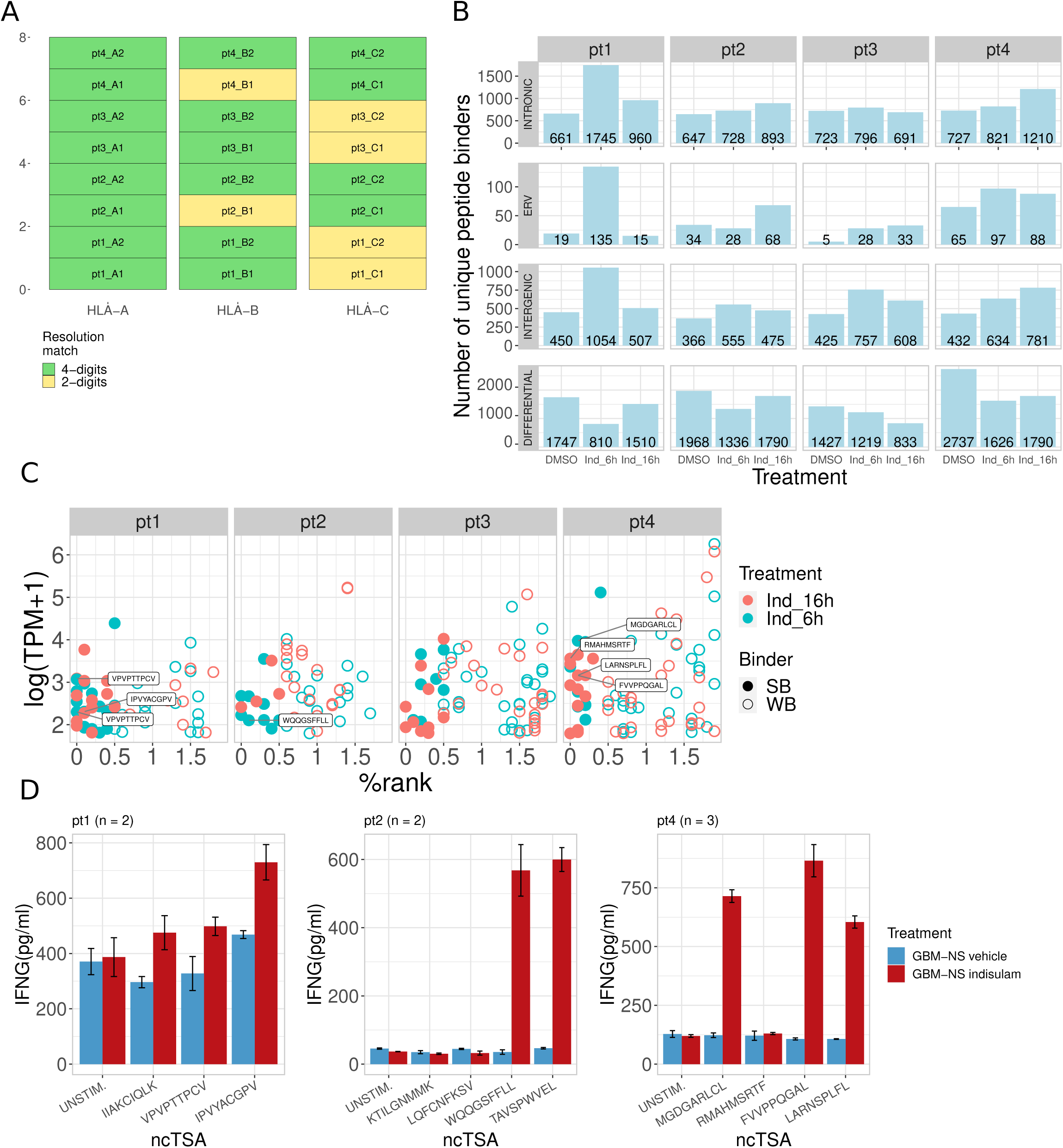
NovumRNA analysis and validation of glioblastoma-derived cell lines treated with indisulam. **A)** Experimental validation of Optitype-based class-I human leukocyte antigen (HLA) typing results. Green fields indicate a match on 4-digits resolution for a specific patient’s sample and HLA allele, while yellow fields indicate an agreement at 2-digit resolution (due to the lower resolution of the polymerase chain reaction (PCR)-based HLA typing). **B)** Bar plots showing the number of different classes of non-canonical tumor specific antigens (ncTSAs) (unique peptide binders) predicted by NovumRNA for the four patients’ cell lines, treated with indisulam for 6 or 16 hours, or with dimethyl sulfoxid (DMSO), used as control. ncTSAs are split by their genomic origin annotation. **C)** Top ncTSA candidates predicted by NovumRNA for the four glioblatoma patients, with reported binding affinity as % rank (x-axis), expression level as log-scaled transcripts per millions (TPM) (y-axis), and time point (indicated by the dot color). The labels indicate the peptide sequence of the ncTSAs selected for validation. **D)** Bar plots summarizing the results of the T-cell essay testing the immunogenicity of the selected ncTSAs. Interferon gamma (IFN-γ) secretion levels in peripheral blood mononuclear cells (PBMCs) co-cultured with tumor cells treated with indisulam (red) or DMSO (blue). PBMCs were either unstimulated (UNSTIM.) or pre-stimulated with the selected ncTSAs predicted via NovumRNA (labelled in panel C). Error bars represent the standard deviation).

We further performed a series of experiments to validate NovumRNA predictions. First, we validated NovumRNA class-I HLA typing results. Using polymerase chain reaction (PCR), we could experimentally determine the HLA alleles at 4-digit resolution for all patients and genes, except for 2 and 4 HLA-B and HLA-C alleles, respectively, which were identified at 2-digit resolution; all the PCR-determined HLA alleles matched the predictions made by NovumRNA (**Fig. 5A**). To validate the predicted ncTSAs, we focused on a selection of *novel* candidates identified from the indisulam-treated samples and not the controls, and having a higher likelihood of being presented and recognized by T cells (**Fig. 5C** and **Table S6**, details in Methods). For this validation, we disregarded patient 3 due to homozygosity for all the three HLA genes (data not shown) and lower resolution available for HLA typing confirmation. We performed a co-culture experiment with autologous PBMCs, either unstimulated or pre-stimulated with the eleven peptides selected for validation, and GB-NS cells treated with indisulam or DMSO. This experiment resulted in an increased secretion of interferon gamma (IFN-γ), as evaluated by ELISA assay, after 24-hour treatment with indisulam compared to the control for six out of the eleven evaluated peptides. Two peptides from patient 2 increased the average IFN-γ secretion upon treatment by 13- and 16-fold, respectively, while three peptides from patient 4 showed a 5-to-8-fold increase. The higher secretion of IFN-γ was consistent with an increased percentage of CD8^+^ T cells CD45RA/CD62L double negative, indicating an effector memory-like phenotype, confirming their immunogenicity as predicted by NovumRNA (**Fig. 5D, Table S8**).

## Discussion

In this study, we introduced NovumRNA, a computational pipeline for the prediction of human non-canonical tumor-specific antigens (ncTSAs) from tumor RNA-seq data. Recent findings have unveiled non-canonical tumor-specific antigens (ncTSAs) as targets of anticancer immune responses^16–22^. Therefore, the ability to computationally predict these antigens from tumor NGS data could significantly broaden the reach of TSA-based cancer immunotherapies, making them accessible and effective for a larger population of patients.

NovumRNA distinguishes itself from previous approaches for ncTSA prediction (e.g.,^19,23–27,29,30,32^) for being a standalone and fully automated software solution which can analyze raw tumor RNA-seq data without relying on third-party tools for data preprocessing. Thanks to its implementation based on Nextflow DSL2^35^ and Singularity containers^45^, NovumRNA ensures portability, scalability, and reproducibility. NovumRNA predicts peptides derived from a wide range of non-canonical, tumor-specific alterations, such as alternative splicing events, expressed retroviral elements, and non-coding regions, and complements its predictions with extensive metadata for downstream neoepitope prioritization and validation. At present, NovumRNA does not predict peptides derived from gene fusions, a task for which we recommend using dedicated tools like NeoFuse^23^ or pVACfuse^28^. The current version of NovumRNA already takes into consideration single nucleotide polymorphisms (SNPs) supported by the majority of the aligned reads, which can alter the sequence of ncTSAs; future improvements will also support the handling of indels as well as of minor-allele SNPs. While our study focuses on class-I ncTSA prediction, we point out that NovumRNA also predicts class-II ncTSAs, which are crucial for the recognition of tumor cells by CD4^+^ T cells^33,34^.

In silico prediction of TSAs is challenging, even for neoantigens derived from sources like SNV and indels^6,44,60^, for which detailed best practices have been proposed^61,62^. Nevertheless, we could demonstrate the robustness of NovumRNA modules and predictions using a diverse panel of publicly-available and ad-hoc-generated datasets with associated gold standards. In particular, using RNA-seq data from a recent proteogenomic study^19^, we could demonstrate a marked agreement of NovumRNA predictions with the original results and superior performance compared to the competitor tool TSAfinder^25^. Of note, unlike the approach used in the original study, NovumRNA predictions were solely based on RNA-seq data and did not require the integration of additional mass spectrometry measurements. While useful for confirming expressed/presented peptides, mass spectrometry entails additional costs and logistical challenges, is rarely available next to RNA-seq in clinical settings and, most importantly, has low sensitivity.

One key challenge in the prediction of ncTSA uniquely from RNA-seq data is the correct translation of aberrant, novel transcripts, which requires the prediction of the exact reading frame (RF) and open reading frame (ORF). For RF prediction, previous studies considered all possible three-frame translations^19,25^. However, this approach results in a broad number of peptides and likely false positives, which can only be reduced via integration of MS-confirmed sequences –performed in the Laumont study^19^ but not by TSAfinder. NovumRNA addresses this by applying a primary filtering step already at the nucleotide level, using reference proteins for translation wherever possible and providing the nucleotide sequence of ncTSAs in the output for further frame testing. NovumRNA accurately predicted 96% of all MS-confirmed peptides reported in the Laumont study, but still obtained diverging results in terms of ncTSA due to alternatively-called RF and ORF. While our test using BORF^53^, a recently-developed tool for ORF prediction, did not result in a marked improvement, we expect that more accurate tools emerging in the near future can be integrated in NovumRNA to further improve its predictions.

False positive predictions, i.e. putative TSAs which are in reality self-peptides, represent another key challenge in ncTSA prediction performed from tumor RNA-seq data alone. NovumRNA reduces false positive rates by leveraging a collection of non-cancerous transcript sequences. The pipeline includes an internal control database derived from TEC RNA-seq data, inspired by the Laumont study^19^, but further allows to flexibly derive larger and/or tissue-specific control databases from user-specified RNA-seq data. We demonstrated the value of this approach by using normal colon RNA-seq data from the GTEx project^54^ to ensure the tumor-specificity of ncTSA candidates predicted from CRC organoid data. Even with only 30 GTEx samples, self-peptide candidates were reduced by 43%, demonstrating the effectiveness of even small collections of non-cancerous samples from the same tissue context. The healthy colon and brain control databases built in this study are available from “https://zenodo.org/records/13642055/files/NovumRNA_resource_bundle.tar.gz?download=1”. Large datasets collections like GTEx or (healthy samples from) TCGA^63^ offer extensive data which can be used to build more extensive, multi-tissue control databases in NovumRNA.

The identification of new immunotherapy targets from non-canonical tumor aberrations can open new therapeutic avenues for patients with hard-to-treat cancers. Low-TMB patients like individuals with MSS CRC are currently not benefitting from cancer immunotherapy, despite responses to ICB immunotherapies have been reported in some settings^14,15^. NovumRNA analysis of RNA-seq data from ten CRC organoids predicted comparable numbers of novel ncTSAs in MSI and MSS tumors, which would suggest similar potential for the presentation of non-canonical tumor-antigen targets. Top candidates ncTSAs identified by NovumRNA, prioritized based on their level of expression and likelihood of being recognized by T cells, included peptides previously associated with melanoma or with the ERVH-5 gene, in line with previous studies^64^. Our analysis further identified a set of putative ncTSAs shared across multiple patient-derived organoids, mainly originating from ERV sequences, which might open the door to “off-the-shelf” therapies.

Glioblastoma represents another hard-to-treat cancer. This extremely aggressive form of brain tumor has been, so far, intractable with both conventional and immunotherapy approaches due to its lowly-immunogenic nature and highly-immunosuppressive milieu^65,66^. Nevertheless, it was reported that patients with high-TMB gliomas, in a context of constitutional DNA mismatch repair deficiency syndrome, can benefit from ICB immunotherapy^67,68^. More recently, it was also shown that recurrent glioblastomas carrying low TMB and an enriched inflammatory gene signature can benefit from immunotherapy and survive longer than recurrent glioblastomas with higher TMB^68^. Motivated by the ncTSA dynamics predicted with NovumRNA on public data from tumor cell lines treated with indisulam, we performed an experiment in glioblastoma primary cell lines perturbed with this splicing-modulating drug. Our investigation included two cell lines derived from recurrent glioblastomas (patient 2 and patient 4) and two from primary glioblastomas. Notably, patient 1 was characterized by an *MGMT*-methylated, hypermethylated, and potentially hypermutated glioblastoma, and previosly showed long-term response to dendritic cell (DC) immunotherapy in the DENDR1 trial (NCT04801147)^69,70^. From this perturbation RNA-seq data, NovumRNA predicted an increase in ncTSAs upon indisulam treatment, with dynamics that were patient-specific. We performed experimental validation of a selection of indisulam-induced ncTSAs, confirming the immunogenicity of 6 out of 11 peptides (54%) specifically in the indisulam-treated samples. The patients’ characteristics summarized above can support the high percentage of immunogenic neoantigens predicted with NovumRNA in our experiment. Our data, together with previous observations describing the ability of indisulam in inducing a proinflammatory microenvironment and increasing CD8^+^ T-cell infiltration^59,71^, support the possibility of expanding the repertoire of therapeutically relevant HLA I-restricted neoepitopes, offering new avenues for low-TMB and antigenically heterogeneous tumors such as glioblastomas. More broadly, these results suggest that NovumRNA can be systematically applied to perturbed tumor RNA-seq data to pinpoint therapeutic interventions that might boost the immunogenicity of tumor cells and facilitate T-cell recognition.

In conclusion, NovumRNA is a comprehensive and fully automated pipeline for the prediction of different classes of ncTSAs from patients’ tumor RNA-seq data. In this study, we demonstrated the value of NovumRNA for pinpointing new candidate targets for immunotherapy and therapeutic interventions which could synergize with anticancer (immuno)therapies, facilitating T-cell mediated recognition of tumor cells. The application of NovumRNA to large cohorts of patients with different cancer types can provide insights on the non-canonical landscape of tumor-specific antigens, which could ultimately guide the design of therapies with higher clinical efficacy and scope.

## Acknowledgments

The computational results presented here have been achieved in part using the LEO HPC infrastructure of the University of Innsbruck. The results shown here are in part based upon data generated by TCGA Research Network (https://cancergenome.nih.gov). The authors would like to thank Anne Krogsdam and the MultiOmics core facility for the support with the tumor organoid sequencing, Benedetta Mazzi (Immuno-hematology and transfusion medicine Unit, San Raffaele Institute) for HLA Typing, Alessandro Gori (SCITEC Institute of Chemical Science and Technology “Giulio Natta”, National Research Council-CNR)for the support with the peptide synthesis.

## STAR Methods

### Lead contact

Further information and requests should be directed to and will be fulfilled by the Lead Contact, Francesca Finotello.

### Materials availability

This study did not generate new unique materials.

### Data and code availability

NovumRNA is available at https://github.com/ComputationalBiomedicineGroup/NovumRNA. RNA-seq data from colorectal cancer organoids and glioblastoma cell lines will be made available through The European Genome-phenome Archive (EGA) upon publication.

## METHOD DETAILS

### The NovumRNA pipeline

NovumRNA is a fully-automated bioinformatic pipeline designed to predict non-canonical tumor-specific antigens (ncTSAs) from raw RNA-seq data. Implemented in the workflow language Nextflow DSL2^35^, NovumRNA ensures ease of use, maximum reproducibility, portability, and parallelism. The pipeline utilizes Singularity containers^45^, managed by Nextflow, which eliminates the need for users to install multiple tools and their dependencies manually.

The pipeline takes as input FASTQ files (zipped or unzipped, single-end or paired-end) from human tumor RNA-seq data. The input data must be provided in a comma-separated values (CSV) formatted sample sheet, containing a unique sample identifier (ID), the sequencing read files (Read1, Read2), and optionally, files containing class-I (HLA_types) and/or class-II HLA types (HLA_types_II). The sample sheet, as well as other parameters, are specified in the novumRNA config file (novumRNA.config). For a detailed description on how to install and use the pipeline, we refer to the NovumRNA documentation on the GitHub repository. Due to licensing restrictions some tools, i.e. netMHCpan and netMHCIIpan^42^, are not shipped with NovumRNA, but they are downloaded from IEDB and installed on the first run after the user accepts the licenses (--accept_license).

Raw sequencing reads are then aligned to the GENCODE reference genome (GRCh38)^39^ using Hisat2^47^, with the option to select STAR^37^ as the aligner. NovumRNA comes with a pre-built HISAT2 index but can also build indices for the specified reference genome and aligner on the fly. StringTie^38^ is used to perform reference-guided transcript assembly (-G gencode.gtf) based on the aligned reads. A custom python script adds the sample name to the Transcript ID in the resulting GTF file and calculates novel features based on the StringTie reported transcript information: isoform_count (the number of transcript isoforms, counted via the assigned StringTie IDs, e.g., STRG.1.1, STRG.1.2). The TPM_iso_perc, the percentage of a transcript’s expression (TPM), relative to the sum of the expression of all isoforms. A high percentage indicates a prevalent abbreviated transcript compared to non abbreviated isoforms, which is crucial to select ncTSAs with higher presentation potential. Exon_cov_ratio, the mean transcript read coverage, divided by the read coverage of each individual exon, to prioritize ncTSAs derived from exons similarly supported by reads like the non abbreviated ones.

NovumRNA employs an innovative filtering strategy based on a so-called capture BED file, to capture novel and differentially expressed transcript fragments from tumor samples, relative to RNA-seq data from healthy tissues. For this, healthy tissue RNA-seq files are aligned to the same reference, and the transcripts reconstructed with StringTie (same as tumor), the resulting GTF files are concatenated to a database which is used to build the capture BED file. The capture BED file consists of two parts: putative novel transcript regions and putative differential regions. Putative novel transcript regions are created by inverting coordinates from the healthy database GTF to identify regions with zero coverage in healthy samples. Based on the coordinates, they are labeled intronic (between exons) or intergenic (between genes). Putative differential regions are regions that are sparsely covered in the healthy database. Bedtools^72^ is used to merge overlapping exon regions from different transcripts in the healthy database GTF, the maximum TPM and maximum Coverage from the exons merged together, is calculated. By default, regions are considered sparse if the maximum TPM < 1 and maximum Coverage < 4, these cut-offs can be changed by the user. NovumRNA arrives with an already built default capture BED file based on 32 RNA-seq samples from human thymic epithelial cells (TECs) originally published by Laumont et al.^19^ and retrieved from the NCBI Gene Expression Omnibus (GEO) with accession codes GSE127825 and GSE127826. This default capture BED file serves as a ground filtering reference, however, users can build their own file based on available data from healthy tissue. To capture the putative regions, the cancer sample GTF file from StringTie is intersected with the capture BED file using bedtools. The captured regions are filtered using a custom python script, keeping only regions longer than 23 nucleotides and from transcripts expressed with at least 1 TPM and covered by >= 4 reads for novel ones, and >=10 and >=16 respectively for differential ones, once more, these cut-offs can be changed by the user.

Transcript fragments are then translated into peptides in two steps: translation of the whole transcript based on the StringTie GTF and translation of tumor-specific regions. For full transcript translation, if a known reference protein is available (“gencode.v41”), the transcript is translated in all three reading frames using a custom Python script and the Biopython^73^ package. The frame that has the longest overlap with the reference protein is selected and trimmed at the first stop codon. If no reference protein is available, the most likely open reading frame is chosen either based on the presence of a Kozak motif, or the closest Methionine to the 5’ end. Cancer-specific regions from the overlap BED file are converted to nucleotide FASTA files using GFFread^74^ and the GENCODE reference genome (GRCh38 version), three-frame translated, and compared to the complete translated transcript. The region showing the highest overlap (minimum of 8 amino acids) is selected and extended to include junction-spanning peptides. A sliding window approach fragments the translated regions into peptides of a specified length, keeping track of the corresponding genomic locations noted in the BED file.

The read coverage of the peptide regions in the original BAM file is checked as an additional quality check, with the BAM file subsetted to retain only high-quality reads. Read coverage is obtained using sam4weblogo.jar^75^, which outputs patient-specific reads covering the regions, considering only reads fully spanning the peptide region. Regions with high coverage are capped at 100 reads, using ‘sortsamrefname.jar’ and ‘biostar154220.jar’^75^, to expedite analysis, and relevant regions are filtered with a custom python script, keeping peptides covered by at least two high-quality reads (cut-off can be changed by user). Simultaneously, patient-specific SNPs are incorporated into the peptides, previously based only on the reference genome sequence. If the total read count and individual count differ, a SNP is present, and the sequence constituting the majority is selected. The selected nucleotide sequences are once more translated to reflect the SNPs at the peptide level.

NovumRNA also performs filtering on a peptide level. The reference proteome (“gencode.v41”) is split into a FASTA file containing all possible peptides of a specified length using a sliding window approach within a custom python script. Only the novel predicted ncTSAs are filtered against this library of healthy reference peptides. The pipeline comes with a default file containing peptides of 9 and 15 amino acids in length, other lengths are built on the fly, if desired.

In parallel, NovumRNA predicts class I HLA types using OptiType^40^. Input RNA-seq FASTQ files are mapped using the YARA mapper^76^ to an HLA RNA reference. The resulting BAM file is used as input for OptiType, which is run in default mode with the –rna option. NovumRNA allows users to skip this step by submitting class I HLA types directly as input within the CSV sample sheet. The HLA typing tool HLA-HD^41^, used for predicting class II HLA types, is not pre-installed with NovumRNA due to licensing issues and must be installed separately by the user, although it is not mandatory for a successful run. To bypass HLA-HD installation, class II HLA types can be directly added to the input CSV samplesheet.

Inferred HLA types are used to predict the binding %rank of the final ncTSAs surviving all filtering steps using netMHCpan or netMHCIIpan^42^, part of the IEDB toolkit^77^ utilized by pVACtools^28^. Only binding peptides are kept, %rank < 2 are labeled as weak binders (WB) and %rank < 0.5 as strong binders (SB).

Finally, all information and generated metadata are consolidated into the final output TSV tables, including additional annotations —such as intronic, intergenic, and differential— based on a BED file containing known ERV regions in the human genome from HERVd^43^. NovumRNA also generates a CSV input sample sheet containing result file paths of already run modules, allowing for reruns without repeating resource-intensive steps like alignment.

### HLA typing benchmark

A gold standard was constructed by leveraging data from two studies that conducted high-precision genotyping of class I (HLA-A, HLA-B, and HLA-C) and class II (HLA-DRB1, HLA-DQB1) HLA types in individuals participating in the 1000 Genomes Project^48–50^, has been done previously^24^ . RNA-seq data of 247 individuals, consensually called in both studies, was retrieved from ArrayExpress (https://www.ebi.ac.uk/arrayexpress, accession: E-GEUV-1). We selected only individuals having calls for every HLA gene in both studies and, after conversion of all HLA types to four-digit resolution, we defined the final consensus types as the intersection of the HLA types reported by both studies for each individual and HLA gene. Each of the tools inferred HLA allele was compared to the gold standard to identify the number of correct (“match”) and wrong (“mismatch”) calls, as well as percentage with respect to the total possible calls. Comparison statistics and plots for this and all other analyses presented in this article were made with custom R scripts, using R version 4.2.3 and ggplot2 v. 3.4.2.

### Assessment of exon-inclusion detection and quantification

Raw human RNA-seq data generated in the rMATS study^52^ was obtained from the NCBI Sequence Read Archive (SRA), under accession code SRS354082. In total, six RNA-seq samples, from two cell lines (PC3E, GS689) with three replicates each, were obtained. Reverse transcription polymerase chain reaction (RT-PCR) validated exon inclusion events, percent spliced in (PSI) values and delta PSI (dPSI) values, i.e. the difference of exon inclusion between two samples/conditions, were obtained from the supplementary material of the original publication. Events genomic coordinates were lifted from the hg19 annotation to the hg38 annotation, using the ‘hgLiftOver’ online tool (https://genome.ucsc.edu/cgi-bin/hgLiftOver).

Events were stored in the BED file format and intersected with the StringTie^38^ assembled GTF derived from NovumRNA analysis of the cell lines, using bedtools intersect^72^. A custom python script was used to identify transcripts including the exon (pre_exon, exon, post_exon) and transcripts excluding the exon (pre_exon, post_exon). Excluding transcripts were defined as to specifically show the pre_exon, followed by the post_exon, possible further isoforms are summed up as “others”. For all transcripts including or excluding the exon, the “TPM_iso_perc” was calculated, i.e. the percentage of the transcript expression in TPM, relative to the sum of the expression in TPM of all transcript isoforms (inclusion + exclusion + others). If multiple transcript isoforms include/exclude the exon, their fractions were summed up. The mean fraction of exon inclusion/exclusion across the three replicates was calculated.

The delta TPM_iso_perc (“dTPM_iso_perc”) was calculated in concordance to the delta PSI (dPSI) values reported in the study (dPSI = PSI_PC3E - PSI_GS689), based on RT-PCR. Pearson correlation was calculated between the average dPSI from the study and NovumRNA’s average “dTPM_iso_perc” per reported exon.

### Validation of NovumRNA using publicly available data

Raw RNA-seq data generated in the Laumont et al. study^19^ from seven primary human cancer samples was retrieved from the gene expression omnibus (GEO) with accession code GSE113972. Mass spectrometry eluted peptides, list of TSAs, healthy control expression, called HLA types and TSAs genomic locations were retrieved from the supplementary materials of the original study.

TSAFinder results obtained on the same dataset were obtained from the supplementary material of TSAFinder publication^25^.

NovumRNA analysis was performed using default parameters, considering peptide lengths of 8, 9, 10, and 11 amino acids.

### Colorectal cancer organoid generation and RNA sequencing

Histologically verified primary colorectal tumor tissues were obtained from patients undergoing surgical resection at the Medical University Hospital of Innsbruck with the approval of the medical ethical committee of the Medical University of Innsbruck for the establishment of colorectal cancer organoid cultures, protocol AN2016-0194 366/4.9. Written informed consent was obtained from the patients prior to surgical sampling. Samples were obtained from adult female or male patients who were treatment-naïve, with the exception of patient CRC39 who received FOLFOX and cetuximab before surgery. Details regarding the patient’s clinical information are provided in **Table S1**.

Colorectal cancer organoids were generated and cultured as previously described^78^. Briefly, freshly isolated tumor cells were seeded in 30µL droplets of 70% Geltrex (ThermoScientific, Cat#A1413202) on 6 wells-plates (Sarstedt, #83.3920), 4 drops each well, in PDO culture medium: Advanced DMEM/F12 (Thermo Scientific, Cat#12634028) supplemented with 10 mM HEPES solution (Sigma, Cat#H0887), 10 mL/L Penicillin-Streptomycin solution, 2 mM GlutaMAX (Thermo Scientific, Cat#3550061), 20% Rspondin conditioned medium, 10% Noggin conditioned medium, 20 mL/L B-27 supplement (Thermo Scientific, Cat#17504044), 1.25 mM N-Acetylcysteine (Sigma, Cat#A9165), 0.5 nM A83-01 (Tocris, Cat#2939), 10 mM SB202190 (Sigma, Cat#S7067), 50 ng/mL human EGF (Peprotech, Cat#AF-100-15), 100 mg/mL Primocin (Invivogen, Cat#ant-pm-2), and 10 mM Y27632 (AbMole, Cat#M1817). PDO culture medium was replaced every two days. To passage the PDOs, Geltrex droplets were dissociated in TripLE Express (Thermo Scientific, Cat#12604013) and re-plated in 30 µL droplets typically with a 1 to 4 split density. PDOs were harvested, snap-frozen and total RNA was isolated using the Nucleospin Mini Kit (Macherey-Nagel, #11912512) following manufacturer’s instruction and submitted to total transcriptome full-length mRNA sequencing (Genewiz for Azenta Life Sciences, Leipzig, Germany).

### Analysis of RNA sequencing data from colorectal cancer organoids

Human healthy colon RNA-seq samples were obtained from the GTEx repository^54^ for filtering. GTEx offers in total 686 healthy colon samples, however, the Bgee suite re-annotated all samples available on GTEx and discarded the ones they deemed unhealthy based on the pathological reports, creating a high-quality subset, restricted to only healthy and non-contaminated samples^79^. Thereby, the total number of purely healthy colon samples we considered for filtering was 262 (list of sample accession identifiers in **Table S2**). FASTQ files underwent preprocessing via NovumRNA, employing the “capture_bed” entry to generate corresponding GTF files. Subsequently, input_samplesheets were constructed, encompassing randomly selected GTF files. This selection process began with 10 files and iteratively increased in increments of 10, with 10 replicates for each step, ultimately yielding 260 distinct samplesheets. Each of these samplesheets served as input for NovumRNA, utilizing the “capture_bed_short” entry to produce 260 corresponding capture BED files. The resulting capture BED files were employed as filtration criteria in NovumRNA. The input data comprised RNA-seq data obtained from 10 colorectal cancer organoids. We applied NovumRNA with default parameters, considering peptides of 9 amino acid length. Jacquard similarity was calculated as the ratio between the number of ncTSAs in the replicates over the number of unique ncTSAs across all replicates.

For each of the ten CRC organoids and each GTEx filtering step (ranging from 10 to 260 samples with increments of ten), as well as for the ten replicates performed at each step, we identified putative self-peptides. These were defined as ncTSAs that were not present, i.e. filtered out, when using the full GTEx dataset (262 samples) for filtering. Using the filtering step with 10 GTEx samples as the reference, we calculated the percentage of removed self-peptides at each filtering step, averaging the result across all replicates and organoids.

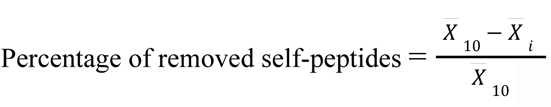, where:

*X̅*_10_ = Average number of self-peptides across all organoids and all replicates at filtering with 10 samples.

*X̅*_*i*_ = Average number of self-peptides across all organoids and all replicates at filtering with *i* samples

(for *i* = 10,…,260).

A Wilcoxon rank-sum test (Mann-Whitney U test) was performed to test the number of predicted ncTSAs between MSI and MSS CRC samples.

Top ncTSA candidates across all organoids at maximum filtering (262 GTEx samples) were selected as only novel (from intronic, or intergenic source), strong- or weak binder to at least one respective HLA type, with a TPM_iso_perc >= 50%, TPM > 3, BAM_reads > 5, and Exon_cov_ratio >= 0.8. From these, only one ncTSA per transcript with the lowest (strongest) binding affinity was chosen.

Seven public antigen databases^55–57,80–82^, (https://caped.icp.ucl.ac.be/), were exported to text files. ncTSAs from 10 CRC organoids, filtered with 262 GTEx samples, were queried in the exported files. ncTSAs needed to be contained in the peptide sequence reported by the database, no mismatch was tolerated.

Predicted ERV derived ncTSA nucleotide sequences, taken from output column “REF_NT”, were used as query and searched in a related nucleotide sequence for gene ERVH-5, taken from AL138920.11 (50173..52469), as reported here https://www.ncbi.nlm.nih.gov/gene/100862699. The search was performed using the SeqKit tool^83^ with the ‘locatè command.

The colon capture_BED file is available as part of the NovumRNA resource bundle: “https://zenodo.org/records/13642055/files/NovumRNA_resource_bundle.tar.gz?download=1”.

### Analysis of public RNA sequencing data from treated cancer cell lines

Raw RNA-seq data from human cancer cell lines that had been subjected to treatments influencing gene transcription were retrieved from two independent studies from the GEO database with accession code GSE162818, and the NCBI SRA database with accession code SRP063667. Subsequently, NovumRNA was employed to analyze each individual sample using default parameters, considering peptides of length 9 amino acids. A one-sided Wilcoxon rank-sum test (Mann-Whitney U test) was performed, using alternative = “greater”, to test if the number of predicted ncTSAs of treated (indisulam, MS023) is greater than the control (DMSO).

### Glioblastoma patients’ characteristics

Glioblastoma samples were provided from patients operated at Fondazione IRCCS Istituto Neurologico Carlo Besta. Written informed consent was obtained from all participants. Clinical data, including age, gender, IDH1 status, and MGMT status are reported in **Table S7** MGMT promoter methylation status was assessed by methylation-specific polymerase chain reaction using specific primers^84^. IDH1 gene mutation detection was performed using primer pairs specific for exon 4 and determined by Sanger sequencing (Applied Biosystems 3500 and 3500 Dx Series Genetic Analyzer). HLA typing was performed using polymerase chain reaction (PCR) using sequence-specific oligonucleotides (PCR-SSO) at Immuno-hematology and transfusion medicine Unit (San Raffaele Institute) on DNA extracted from blood samples (Puregene Blood Core Kit, Qiagen).

### Generation of RNA sequencing data from glioblastoma primary cell lines treated with indisulam

Four glioblastoma primary cell lines ( BT592/pt1, BT1007/pt2, BT1009/pt3; BT1012/pt4) were derived from glioblastoma Cavitron Ultrasonic Aspirator (CUSA) material as previously described^85,86^ and cultured as neurospheres (GB-NS) in DMEM/F12+Glutamax medium containing B27 supplement (Thermofisher, Waltham, MA, USA) and the mitogenic factors epidermal growth factor (EGF) and fibroblast growth factor b (bFGF) (Peprotech, Rocky Hill, NJ, USA). GB-NS were confirmed mycoplasma-free by PCR Test. Cells were seeded in triplicate in 6-well plates, treated with 5 μM indisulam (Sigma Aldrich, Merk), or vehicle control, and collected after 6 and 16 hours for RNA extraction.

Total RNA was extracted using TRIzol Reagent (Life Technologies, Thermo Fisher, cat n. 15596026), RNeasy Mini Kit, and RNase-Free DNase Set (Qiagen, cat n. 74104 and 79254). Sample QC, RNA library preparations, and sequencing reactions were conducted at GENEWIZ (Germany GmbH). The concentration of RNA was quantified using a Qubit Fluorometer (Life Technologies, Carlsbad, CA) and RNA integrity was assayed using a TapeStation. Sequencing libraries were prepared with RNA with PolyA selection and ERCC spike-in with standard ERCC kit (ERCC RNA Spike-In Mix,Thermo Fisher 4456740). Final libraries (Illumina, RNA with PolyA selection) were sequenced on a NovaSeq platform, producing 2×150 base paired-end reads.

### Analysis of RNA sequencing data from glioblastoma cell lines treated with indisulam and selection of top candidates for validation

NovumRNA was employed to analyze each individual sample using a capture_BED file built based on 120 healthy brain GTEx samples deemed purely healthy by the bgee suite^79^ (list of sample accession identifiers in **Table S2**), predicting peptides of 9 amino acids length.

Predicted peptides submitted for validation from patients 1, 2, and 4, were selected based on desirable metadata features: Predicted only after indisulam treatment, so not found in the DMSO control, only strong binders (%rank < 0.5), TPM_iso_perc = 100, Exon_cov_ratio >= 0.8 and TPM >= 0.5, BAM_reads >= 5. From the remaining pool of peptides fulfilling these features, only one peptide per transcript was selected based on the lowest %rank score. From these, four peptides per patient with the lowest %rank were chosen. For patient 1, a peptide was selected which occurred with six hours and sixteen hours indisulam treatment, leading to three individual peptides validated.

The brain capture_BED file is available as part of the NovumRNA resource bundle: “https://zenodo.org/records/13642055/files/NovumRNA_resource_bundle.tar.gz?download=1”.

### Experimental validation via co-culture experiments

A total of 2×10^5^ autologous dendritic cells (DCs) were plated in 24 well-plates, pre-loaded with 10 microg/ml of each selected peptide (synthetized by Primm) for 2 hours at 37°C and then co-cultured with 2×10^6^ PBMCs (ratio 1:10 = DC:PBMCs) for three days in a final volume of 2 ml RPMI 1640 supplemented with 10% human serum, 100 U/ml penicillin, 100 U/ml streptomycin, 100 µg/ml glutamine, 0.1 mM non-essential amino acids, 1 mM sodium pyruvate, 50 µM β-mercaptoethanol the presence of 30 U/ml of IL-2 (Miltenyi Biotec, Germany).

Pre-stimulated PBMCs were co-cultured in the presence of GB-NS properly treated with indisulam or DMSO as control (E:T = 3:1) in GB-NS medium without serum and in the presence of B27 supplement. Co-culture supernatants were collected after 24 hours to measure IFN-γ using specific ELISA (R&D system, Minneapolis, MN). The memory status of T cells was assessed by using CD45RA and CD62L that distinguish stem memory/naïve (CD45RA^+^ CD62L^+^); effector memory (CD45RA^−^ CD62L^−^); central memory (CD45RA− CD62L^+^).

A one-sided Wilcoxon rank-sum test (Mann-Whitney U test) was performed, using alternative = “greater”, to test for increased secretion of IFN-γ for pre-stimulated PBMCs and GB-NS cells treated with indisulam or DMSO, compared to the control for evaluated peptides.

## Supplemental information

**Document_S1**. Figures S1–S3 and Tables S1,S7,S8

**Document_S2**. Excel file containing Tables S2-S6

## References

1. Galluzzi, L., Vacchelli, E., Bravo-San Pedro, J.-M., Buqué, A., Senovilla, L., Baracco, E.E., Bloy, N., Castoldi, F., Abastado, J.-P., Agostinis, P., et al. (2014). Classification of current anticancer immunotherapies. Oncotarget 5, 12472–12508.

2. Schumacher, T.N., and Schreiber, R.D. (2015). Neoantigens in cancer immunotherapy. Science 348, 69–74.

3. Schumacher, T.N., Scheper, W., and Kvistborg, P. (2019). Cancer Neoantigens. Annu. Rev. Immunol. 37, 173–200.

4. Hackl, H., Charoentong, P., Finotello, F., and Trajanoski, Z. (2016). Computational genomics tools for dissecting tumour-immune cell interactions. Nat. Rev. Genet. 17, 441–458.

5. Chen, D.S., and Mellman, I. (2013). Oncology meets immunology: the cancer-immunity cycle. Immunity 39, 1–10.

6. Lee, C.-H., Yelensky, R., Jooss, K., and Chan, T.A. (2018). Update on Tumor Neoantigens and Their Utility: Why It Is Good to Be Different. Trends Immunol. 39, 536–548.

7. Havel, J.J., Chowell, D., and Chan, T.A. (2019). The evolving landscape of biomarkers for checkpoint inhibitor immunotherapy. Nat. Rev. Cancer 19, 133–150.

8. Haslam, A., and Prasad, V. (2019). Estimation of the Percentage of US Patients With Cancer Who Are Eligible for and Respond to Checkpoint Inhibitor Immunotherapy Drugs. JAMA Netw Open 2, e192535.

9. Pilard, C., Ancion, M., Delvenne, P., Jerusalem, G., Hubert, P., and Herfs, M. (2021). Cancer immunotherapy: it’s time to better predict patients’ response. Br. J. Cancer. 10.1038/s41416-021-01413-x.

10. Le, D.T., Durham, J.N., Smith, K.N., Wang, H., Bartlett, B.R., Aulakh, L.K., Lu, S., Kemberling, H., Wilt, C., Luber, B.S., et al. (2017). Mismatch repair deficiency predicts response of solid tumors to PD-1 blockade. Science 357, 409–413.

11. Asaoka, Y., Ijichi, H., and Koike, K. (2015). PD-1 Blockade in Tumors with Mismatch-Repair Deficiency. N. Engl. J. Med. 373, 1979.

12. Overman, M.J., McDermott, R., Leach, J.L., Lonardi, S., Lenz, H.-J., Morse, M.A., Desai, J., Hill, A., Axelson, M., Moss, R.A., et al. (2017). Nivolumab in patients with metastatic DNA mismatch repair-deficient or microsatellite instability-high colorectal cancer (CheckMate 142): an open-label, multicentre, phase 2 study. Lancet Oncol. 18, 1182–1191.

13. Litchfield, K., Reading, J.L., Puttick, C., Thakkar, K., Abbosh, C., Bentham, R., Watkins, T.B.K., Rosenthal, R., Biswas, D., Rowan, A., et al. (2021). Meta-analysis of tumor- and T cell-intrinsic mechanisms of sensitization to checkpoint inhibition. Cell 184, 596–614.e14.

14. Yarchoan, M., Hopkins, A., and Jaffee, E.M. (2017). Tumor Mutational Burden and Response Rate to PD-1 Inhibition. N. Engl. J. Med. 377, 2500–2501.

15. Chalabi, M., Fanchi, L.F., Dijkstra, K.K., Van den Berg, J.G., Aalbers, A.G., Sikorska, K., Lopez-Yurda, M., Grootscholten, C., Beets, G.L., Snaebjornsson, P., et al. (2020). Neoadjuvant immunotherapy leads to pathological responses in MMR-proficient and MMR-deficient early-stage colon cancers. Nat. Med. 26, 566–576.

16. Kahles, A., Lehmann, K.-V., Toussaint, N.C., Hüser, M., Stark, S.G., Sachsenberg, T., Stegle, O., Kohlbacher, O., Sander, C., Cancer Genome Atlas Research Network, et al. (2018). Comprehensive Analysis of Alternative Splicing Across Tumors from 8,705 Patients. Cancer Cell 34, 211–224.e6.

17. Kumar-Sinha, C., Kalyana-Sundaram, S., and Chinnaiyan, A.M. (2015). Landscape of gene fusions in epithelial cancers: seq and ye shall find. Genome Med. 7, 129.

18. Attig, J., Young, G.R., Hosie, L., Perkins, D., Encheva-Yokoya, V., Stoye, J.P., Snijders, A.P., Ternette, N., and Kassiotis, G. (2019). LTR retroelement expansion of the human cancer transcriptome and immunopeptidome revealed by de novo transcript assembly. Genome Res. 10.1101/gr.248922.119.

19. Laumont, C.M., Vincent, K., Hesnard, L., Audemard, É., Bonneil, É., Laverdure, J.-P., Gendron, P., Courcelles, M., Hardy, M.-P., Côté, C., et al. (2018). Noncoding regions are the main source of targetable tumor-specific antigens. Sci. Transl. Med. 10. 10.1126/scitranslmed.aau5516.

20. Xiang, R., Ma, L., Yang, M., Zheng, Z., Chen, X., Jia, F., Xie, F., Zhou, Y., Li, F., Wu, K., et al. (2021). Increased expression of peptides from non-coding genes in cancer proteomics datasets suggests potential tumor neoantigens. Commun Biol 4, 496.

21. Smith, C.C., Selitsky, S.R., Chai, S., Armistead, P.M., Vincent, B.G., and Serody, J.S. (2019). Alternative tumour-specific antigens. Nat. Rev. Cancer 19, 465–478.

22. Shi, Y., Jing, B., and Xi, R. (2023). Comprehensive analysis of neoantigens derived from structural variation across whole genomes from 2528 tumors. Genome Biol. 24, 169.

23. Fotakis, G., Rieder, D., Haider, M., Trajanoski, Z., and Finotello, F. (2020). NeoFuse: predicting fusion neoantigens from RNA sequencing data. Bioinformatics 36, 2260–2261.

24. Rieder, D., Fotakis, G., Ausserhofer, M., René, G., Paster, W., Trajanoski, Z., and Finotello, F. (2021). nextNEOpi: a comprehensive pipeline for computational neoantigen prediction. Bioinformatics. 10.1093/bioinformatics/btab759.

25. Sharpnack, M.F., Johnson, T.S., Chalkley, R., Han, Z., Carbone, D., Huang, K., and He, K. (2022). TSAFinder: exhaustive tumor-specific antigen detection with RNAseq. Bioinformatics 38, 2422–2427.

26. Chai, S., Smith, C.C., Kochar, T.K., Hunsucker, S.A., Beck, W., Olsen, K.S., Vensko, S., Glish, G.L., Armistead, P.M., Prins, J.F., et al. (2022). NeoSplice: a bioinformatics method for prediction of splice variant neoantigens. Bioinform Adv 2, vbac032.

27. Lang, F., Sorn, P., Suchan, M., Henrich, A., Albrecht, C., Köhl, N., Beicht, A., Riesgo-Ferreiro, P., Holtsträter, C., Schrörs, B., et al. (2024). Prediction of tumor-specific splicing from somatic mutations as a source of neoantigen candidates. Bioinform Adv 4, vbae080.

28. Hundal, J., Kiwala, S., McMichael, J., Miller, C.A., Xia, H., Wollam, A.T., Liu, C.J., Zhao, S., Feng, Y.-Y., Graubert, A.P., et al. (2020). pVACtools: A Computational Toolkit to Identify and Visualize Cancer Neoantigens. Cancer Immunol Res 8, 409–420.

29. Li, G., Mahajan, S., Ma, S., Jeffery, E.D., Zhang, X., Bhattacharjee, A., Venkatasubramanian, M., Weirauch, M.T., Miraldi, E.R., Grimes, H.L., et al. (2024). Splicing neoantigen discovery with SNAF reveals shared targets for cancer immunotherapy. Sci. Transl. Med. 16, eade2886.

30. Zhang, J., Mardis, E.R., and Maher, C.A. (2017). INTEGRATE-neo: a pipeline for personalized gene fusion neoantigen discovery. Bioinformatics 33, 555–557.

31. Tan, X., Xu, L., Jian, X., Ouyang, J., Hu, B., Yang, X., Wang, T., and Xie, L. (2023). PGNneo: A Proteogenomics-Based Neoantigen Prediction Pipeline in Noncoding Regions. Cells 12. 10.3390/cells12050782.

32. Liu, C., Zhang, Y., Jian, X., Tan, X., Lu, M., Ouyang, J., Liu, Z., Li, Y., Xu, L., Chen, L., et al. (2022). ProGeo-Neo v2.0: A One-Stop Software for Neoantigen Prediction and Filtering Based on the Proteogenomics Strategy. Genes 13. 10.3390/genes13050783.

33. Ahrends, T., and Borst, J. (2018). The opposing roles of CD4+ T cells in anti-tumour immunity. Immunology 154, 582–592.

34. Tay, R.E., Richardson, E.K., and Toh, H.C. (2021). Revisiting the role of CD4+ T cells in cancer immunotherapy-new insights into old paradigms. Cancer Gene Ther. 28, 5–17.

35. Di Tommaso, P., Chatzou, M., Floden, E.W., Barja, P.P., Palumbo, E., and Notredame, C. (2017). Nextflow enables reproducible computational workflows. Nat. Biotechnol. 35, 316–319.

36. Kim, D., Paggi, J.M., Park, C., Bennett, C., and Salzberg, S.L. (2019). Graph-based genome alignment and genotyping with HISAT2 and HISAT-genotype. Nat. Biotechnol. 37, 907–915.

37. Dobin, A., Davis, C.A., Schlesinger, F., Drenkow, J., Zaleski, C., Jha, S., Batut, P., Chaisson, M., and Gingeras, T.R. (2013). STAR: ultrafast universal RNA-seq aligner. Bioinformatics 29, 15–21.

38. Pertea, M., Pertea, G.M., Antonescu, C.M., Chang, T.-C., Mendell, J.T., and Salzberg, S.L. (2015). StringTie enables improved reconstruction of a transcriptome from RNA-seq reads. Nat. Biotechnol. 33, 290–295.

39. Frankish, A., Carbonell-Sala, S., Diekhans, M., Jungreis, I., Loveland, J.E., Mudge, J.M., Sisu, C., Wright, J.C., Arnan, C., Barnes, I., et al. (2023). GENCODE: reference annotation for the human and mouse genomes in 2023. Nucleic Acids Res. 51, D942–D949.

40. Szolek, A., Schubert, B., Mohr, C., Sturm, M., Feldhahn, M., and Kohlbacher, O. (2014). OptiType: precision HLA typing from next-generation sequencing data. Bioinformatics 30, 3310–3316.

41. Kawaguchi, S., Higasa, K., Shimizu, M., Yamada, R., and Matsuda, F. (2017). HLA-HD: An accurate HLA typing algorithm for next-generation sequencing data. Hum. Mutat. 38, 788–797.

42. Reynisson, B., Alvarez, B., Paul, S., Peters, B., and Nielsen, M. (2020). NetMHCpan-4.1 and NetMHCIIpan-4.0: improved predictions of MHC antigen presentation by concurrent motif deconvolution and integration of MS MHC eluted ligand data. Nucleic Acids Res. 48, W449–W454.

43. Paces, J., Pavlícek, A., and Paces, V. (2002). HERVd: database of human endogenous retroviruses. Nucleic Acids Res. 30, 205–206.

44. Wells, D.K., van Buuren, M.M., Dang, K.K., Hubbard-Lucey, V.M., Sheehan, K.C.F., Campbell, K.M., Lamb, A., Ward, J.P., Sidney, J., Blazquez, A.B., et al. (2020). Key Parameters of Tumor Epitope Immunogenicity Revealed Through a Consortium Approach Improve Neoantigen Prediction. Cell 183, 818–834.e13.

45. Kurtzer, G.M., Sochat, V., and Bauer, M.W. (2017). Singularity: Scientific containers for mobility of compute. PLoS One 12, e0177459.

46. Orenbuch, R., Filip, I., Comito, D., Shaman, J., Pe’er, I., and Rabadan, R. (2020). arcasHLA: high-resolution HLA typing from RNAseq. Bioinformatics 36, 33–40.

47. Kim, D., Paggi, J., and Salzberg, S.L. (2018). HISAT-genotype: Next Generation Genomic Analysis Platform on a Personal Computer. bioRxiv, 266197. 10.1101/266197.

48. Gourraud, P.-A., Khankhanian, P., Cereb, N., Yang, S.Y., Feolo, M., Maiers, M., Rioux, J.D., Hauser, S., and Oksenberg, J. (2014). HLA diversity in the 1000 genomes dataset. PLoS One 9, e97282.

49. Abi-Rached, L., Gouret, P., Yeh, J.-H., Di Cristofaro, J., Pontarotti, P., Picard, C., and Paganini, J. (2018). Immune diversity sheds light on missing variation in worldwide genetic diversity panels. PLoS One 13, e0206512.

50. 1000 Genomes Project Consortium, Auton, A., Brooks, L.D., Durbin, R.M., Garrison, E.P., Kang, H.M., Korbel, J.O., Marchini, J.L., McCarthy, S., McVean, G.A., et al. (2015). A global reference for human genetic variation. Nature 526, 68–74.

51. Jiang, M., Zhang, S., Yin, H., Zhuo, Z., and Meng, G. (2023). A comprehensive benchmarking of differential splicing tools for RNA-seq analysis at the event level. Brief. Bioinform. 24. 10.1093/bib/bbad121.

52. Shen, S., Park, J.W., Lu, Z.-X., Lin, L., Henry, M.D., Wu, Y.N., Zhou, Q., and Xing, Y. (2014). rMATS: robust and flexible detection of differential alternative splicing from replicate RNA-Seq data. Proc. Natl. Acad. Sci. U. S. A. 111, E5593–E5601.

53. Signal, B., and Kahlke, T. (2021). Borf: Improved ORF prediction in de-novo assembled transcriptome annotation. bioRxiv, 2021.04.12.439551. 10.1101/2021.04.12.439551.

54. GTEx Consortium, Laboratory, Data Analysis &Coordinating Center (LDACC)—Analysis Working Group, Statistical Methods groups—Analysis Working Group, Enhancing GTEx (eGTEx) groups, NIH Common Fund, NIH/NCI, NIH/NHGRI, NIH/NIMH, NIH/NIDA, Biospecimen Collection Source Site—NDRI, et al. (2017). Genetic effects on gene expression across human tissues. Nature 550, 204–213.

55. Othoum, G., and Maher, C.A. (2023). CrypticProteinDB: an integrated database of proteome and immunopeptidome derived non-canonical cancer proteins. NAR Cancer 5, zcad024.

56. Tan, X., Li, D., Huang, P., Jian, X., Wan, H., Wang, G., Li, Y., Ouyang, J., Lin, Y., and Xie, L. (2020). dbPepNeo: a manually curated database for human tumor neoantigen peptides. Database 2020. 10.1093/database/baaa004.

57. Xia, J., Bai, P., Fan, W., Li, Q., Li, Y., Wang, D., Yin, L., and Zhou, Y. (2021). NEPdb: A Database of T-Cell Experimentally-Validated Neoantigens and Pan-Cancer Predicted Neoepitopes for Cancer Immunotherapy. Front. Immunol. 12, 644637.

58. Ding, X.-L., Yang, X., Liang, G., and Wang, K. (2016). Isoform switching and exon skipping induced by the DNA methylation inhibitor 5-Aza-2’-deoxycytidine. Sci. Rep. 6, 24545.

59. Lu, S.X., De Neef, E., Thomas, J.D., Sabio, E., Rousseau, B., Gigoux, M., Knorr, D.A., Greenbaum, B., Elhanati, Y., Hogg, S.J., et al. (2021). Pharmacologic modulation of RNA splicing enhances anti-tumor immunity. Cell 184, 4032–4047.e31.

60. Saini, S.K., Rekers, N., and Hadrup, S.R. (2017). Novel tools to assist neoepitope targeting in personalized cancer immunotherapy. Ann. Oncol. 28, xii3–xii10.

61. De Mattos-Arruda, L., Vazquez, M., Finotello, F., Lepore, R., Porta, E., Hundal, J., Amengual-Rigo, P., Ng, C.K.Y., Valencia, A., Carrillo, J., et al. (2020). Neoantigen prediction and computational perspectives towards clinical benefit: recommendations from the ESMO Precision Medicine Working Group. Ann. Oncol. 31, 978–990.

62. Richters, M.M., Xia, H., Campbell, K.M., Gillanders, W.E., Griffith, O.L., and Griffith, M. (2019). Best practices for bioinformatic characterization of neoantigens for clinical utility. Genome Med. 11, 56.

63. Cancer Genome Atlas Research Network, Weinstein, J.N., Collisson, E.A., Mills, G.B., Shaw, K.R.M., Ozenberger, B.A., Ellrott, K., Shmulevich, I., Sander, C., and Stuart, J.M. (2013). The Cancer Genome Atlas Pan-Cancer analysis project. Nat. Genet. 45, 1113–1120.

64. Rooney, M.S., Shukla, S.A., Wu, C.J., Getz, G., and Hacohen, N. (2015). Molecular and genetic properties of tumors associated with local immune cytolytic activity. Cell 160, 48–61.

65. Wang, X., Lu, J., Guo, G., and Yu, J. (2021). Immunotherapy for recurrent glioblastoma: practical insights and challenging prospects. Cell Death Dis. 12, 299.

66. Sharma, P., Aaroe, A., Liang, J., and Puduvalli, V.K. (2023). Tumor microenvironment in glioblastoma: Current and emerging concepts. Neurooncol Adv 5, vdad009.

67. Touat, M., Li, Y.Y., Boynton, A.N., Spurr, L.F., Iorgulescu, J.B., Bohrson, C.L., Cortes-Ciriano, I., Birzu, C., Geduldig, J.E., Pelton, K., et al. (2020). Mechanisms and therapeutic implications of hypermutation in gliomas. Nature 580, 517–523.

68. Gromeier, M., Brown, M.C., Zhang, G., Lin, X., Chen, Y., Wei, Z., Beaubier, N., Yan, H., He, Y., Desjardins, A., et al. (2021). Very low mutation burden is a feature of inflamed recurrent glioblastomas responsive to cancer immunotherapy. Nat. Commun. 12, 352.

69. Pellegatta, S., Eoli, M., Cuccarini, V., Anghileri, E., Pollo, B., Pessina, S., Frigerio, S., Servida, M., Cuppini, L., Antozzi, C., et al. (2018). Survival gain in glioblastoma patients treated with dendritic cell immunotherapy is associated with increased NK but not CD8+ T cell activation in the presence of adjuvant temozolomide. Oncoimmunology 7, e1412901.

70. Miele, E., Anghileri, E., Calatozzolo, C., Lazzarini, E., Patrizi, S., Ciolfi, A., Pedace, L., Patanè, M., Abballe, L., Paterra, R., et al. (2024). Clinicopathological and molecular landscape of 5-year IDH-wild-type glioblastoma survivors: A multicentric retrospective study. Cancer Lett. 588, 216711.

71. Ippolito, M.R., Zerbib, J., Eliezer, Y., Reuveni, E., Viganò, S., De Feudis, G., Kadmon, A.S., Vigorito, I., Martin, S., Laue, K., et al. (2023). Increased RNA and protein degradation is required for counteracting transcriptional burden and proteotoxic stress in human aneuploid cells. bioRxiv, 2023.01.27.525826. 10.1101/2023.01.27.525826.

72. Quinlan, A.R., and Hall, I.M. (2010). BEDTools: a flexible suite of utilities for comparing genomic features. Bioinformatics 26, 841–842.

73. Cock, P.J.A., Antao, T., Chang, J.T., Chapman, B.A., Cox, C.J., Dalke, A., Friedberg, I., Hamelryck, T., Kauff, F., Wilczynski, B., et al. (2009). Biopython: freely available Python tools for computational molecular biology and bioinformatics. Bioinformatics 25, 1422–1423.

74. Pertea, G., and Pertea, M. (2020). GFF Utilities: GffRead and GffCompare. F1000Res. 9, 304.

75. Lindenbaum, P. (2015). JVarkit: java-based utilities for Bioinformatics. Preprint at figshare, 10.6084/M9.FIGSHARE.1425030.V1 10.6084/M9.FIGSHARE.1425030.V1.

76. Dadi, T.H., Siragusa, E., Piro, V.C., Andrusch, A., Seiler, E., Renard, B.Y., and Reinert, K. (2018). DREAM-Yara: an exact read mapper for very large databases with short update time. Bioinformatics 34, i766–i772.

77. Vita, R., Mahajan, S., Overton, J.A., Dhanda, S.K., Martini, S., Cantrell, J.R., Wheeler, D.K., Sette, A., and Peters, B. (2019). The Immune Epitope Database (IEDB): 2018 update. Nucleic Acids Res. 47, D339–D343.

78. Plattner, C., Lamberti, G., Blattmann, P., Kirchmair, A., Rieder, D., Loncova, Z., Sturm, G., Scheidl, S., Ijsselsteijn, M., Fotakis, G., et al. (2023). Functional and spatial proteomics profiling reveals intra- and intercellular signaling crosstalk in colorectal cancer. iScience 26, 108399.

79. Bastian, F.B., Roux, J., Niknejad, A., Comte, A., Fonseca Costa, S.S., de Farias, T.M., Moretti, S., Parmentier, G., de Laval, V.R., Rosikiewicz, M., et al. (2021). The Bgee suite: integrated curated expression atlas and comparative transcriptomics in animals. Nucleic Acids Res. 49, D831–D847.

80. Wu, T., Chen, J., Diao, K., Wang, G., Wang, J., Yao, H., and Liu, X.-S. (2023). Neodb: a comprehensive neoantigen database and discovery platform for cancer immunotherapy. Database 2023. 10.1093/database/baad041.

81. Wu, J., Chen, W., Zhou, Y., Chi, Y., Hua, X., Wu, J., Gu, X., Chen, S., and Zhou, Z. (2023). TSNAdb v2.0: The Updated Version of Tumor-specific Neoantigen Database. Genomics Proteomics Bioinformatics 21, 259–266.

82. Yi, X., Liao, Y., Wen, B., Li, K., Dou, Y., Savage, S.R., and Zhang, B. (2021). caAtlas: An immunopeptidome atlas of human cancer. iScience 24, 103107.

83. Shen, W., Le, S., Li, Y., and Hu, F. (2016). SeqKit: A Cross-Platform and Ultrafast Toolkit for FASTA/Q File Manipulation. PLoS One 11, e0163962.

84. Eoli, M., Menghi, F., Bruzzone, M.G., De Simone, T., Valletta, L., Pollo, B., Bissola, L., Silvani, A., Bianchessi, D., D’Incerti, L., et al. (2007). Methylation of O6-methylguanine DNA methyltransferase and loss of heterozygosity on 19q and/or 17p are overlapping features of secondary glioblastomas with prolonged survival. Clin. Cancer Res. 13, 2606–2613.

85. Behnan, J., Stangeland, B., Langella, T., Finocchiaro, G., Murrell, W., and Brinchmann, J.E. (2016). Ultrasonic Surgical Aspirate is a Reliable Source For Culturing Glioblastoma Stem Cells. Sci. Rep. 6, 32788.

86. Finocchiaro, G., and Pellegatta, S. (2016). Immunotherapy with dendritic cells loaded with glioblastoma stem cells: from preclinical to clinical studies. Cancer Immunol. Immunother. 65, 101–109.

