## Supplementary material for "NovumRNA: accurate prediction of non-canonical tumor antigens from RNA sequencing data": Document_S1

### Supplementary tables

**Table S1 | Metadata from 10 CRC organoids.**
MSI: Microsatellite instable (associated with high load of mutations).
MSS: Microsatellite stable.

| ID | Gender | Age | Subtype | Tumor location |
|---|---|---|---|---|
| CRC01 | male | 75 | MSI | ascending colon |
| CRC13 | female | 90 | MSI | sigmoid colon |
| CRC22 | male | 87 | MSI | ascending colon |
| CRC34 | male | 52 | unknown | sigmoid colon |
| CRC35 | female | 74 | MSS | sigmoid colon |
| CRC39 | male | 44 | MSS | sigmoid colon |
| CRC41 | female | 37 | MSS | sigmoid colon |
| CRC42 | male | 60 | MSS | sigmoid colon |
| CRC43 | female | 68 | MSS | cecum |
| CRC44 | male | 58 | MSS | sigmoid colon |

**Table S7 | Metadata from 4 glioblastoma patients.**

| Pt ID | GB-NS ID | Diagnosis | Age at surgery | Gend er | IDH1 | MGMT methylation |
|---|---|---|---|---|---|---|
| Patient 1* | BT592 | glioblastoma | 45 | M | WT | MET (0.74) |
| Patient 2 | BT1007 | recurrent glioblastoma | 59 | M | WT | UNMET (0.00) |
| Patient 3 | BT1009 | glioblastoma | 77 | M | WT | UNMET (0.00) |
| Patient 4 | BT1012 | recurrent glioblastoma | 76 | F | WT | MET (2.48) |

*Patient 1 is Pt23 in DENDR1 clinical trial (NCT04801147).
GB-NS: glioblastoma neurospheres; IDH: Isocitrate dehydrogenase; MGMT: O-6-methylguanine-DNA methyltransferase.

**Table S8 | Memory of T cells after peptide stimulation.**

| Patient | Peptide | Treatment | %CD8+ T stem memory/naive | %CD8+ T central memory | %CD8+ T effector memory |
|---|---|---|---|---|---|
| Patient 1/ BT592 | unstimulated | vehicle | 17.51 | 23.01 | 38.61 |
| | | Indisulam | 19.40 | 30.94 | 36.18 |
| | IIAKCIQLK | vehicle | 21.91 | 46.59 | 29.91 |
| | | Indisulam | 18.02 | 43.81 | 25.59 |
| | VPVPTTPCV | vehicle | 27.80 | 44.94 | 21.95 |
| | | Indisulam | 23.00 | 45.75 | 31.24 |
| | **IPVYACGPV** | **vehicle** | **25.61** | **48.11** | **31.46** |
| | | **Indisulam** | **10.20** | **38.72** | **71.20** |
| Patient 2 / BT1007 | unstimulated | vehicle | 25.6 | 32.4 | 33.8 |
| | | Indisulam | 24.9 | 33.9 | 35.3 |
| | KTILGNMMK | vehicle | 15.4 | 34.7 | 33.2 |
| | | Indisulam | 17.5 | 36.1 | 38.4 |
| | LQFCNFKSV | vehicle | 17.5 | 31.8 | 37.3 |
| | | Indisulam | 14.7 | 32.8 | 36.7 |
| | **WQQGSFFLL** | **vehicle** | **19.9** | **38.2** | **33.9** |
| | | **Indisulam** | **9.8** | **36** | **50.1** |
| | **TAVSPWVEL** | **vehicle** | **28.7** | **28.6** | **41.7** |
| | | **Indisulam** | **10.2** | **39.7** | **51.2** |
| Patient 4 / BT1012 | unstimulated | vehicle | 23.8 | 29.5 | 42.9 |
| | | Indisulam | 22.6 | 33.3 | 42.6 |
| | **MGDGARLCL** | **vehicle** | **26.9** | **32.1** | **22.1** |
| | | **Indisulam** | **13.7** | **22.3** | **55.3** |
| | RMAHMSRTF | vehicle | 20.8 | 44.8 | 31.9 |
| | | Indisulam | 23.3 | 42.7 | 31.2 |
| | **FVVPPQGAL** | **vehicle** | **20.3** | **29.4** | **30.53** |
| | | **Indisulam** | **12.4** | **26.4** | **42.1** |
| | **LARNSPLFL** | **vehicle** | **21.1** | **41.4** | **29.9** |
| | | **Indisulam** | **13.4** | **43.2** | **40.7** |

stem memory/naïve: CD45RA+ CD62L+; effector memory: CD45RA− CD62L−;
central memory: CD45RA− CD62L+

### Supplementary figures

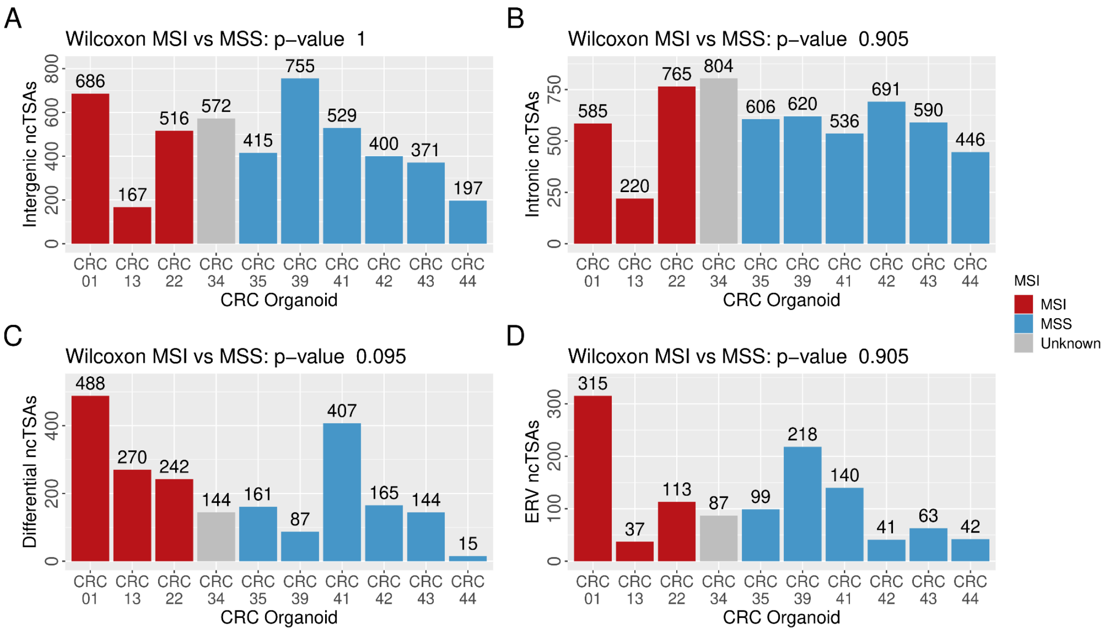


**Figure S1 | NovumRNA predicted ncTSAs in MSI versus MSS CRC organoids, filtered using 262 healthy colon samples from GTEx.** ncTSAs of different origin predicted by NovumRNA per colorectal cancer (CRC) organoid, colored by the organoid subtype. A Wilcoxon test was performed between predictions of MSI and MSS samples.

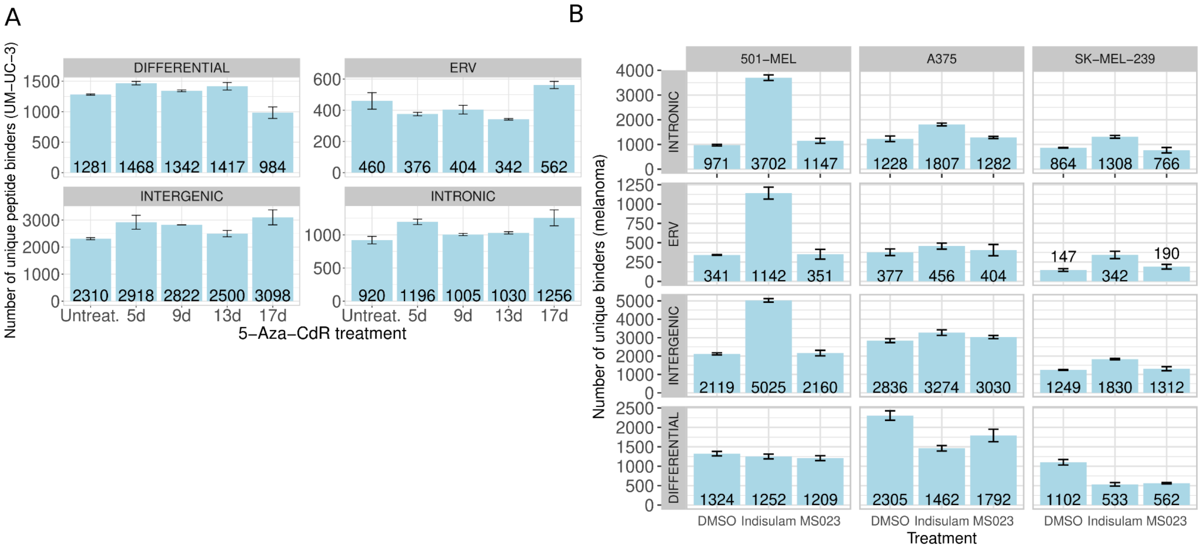


**Figure S2 | Predicted ncTSAs in splicing perturbed human cell lines.**

**A) Predicted ncTSAs in urinary bladder cancer cell lines treated with 5-Aza-CdR.** The bar plot shows the mean number of unique ncTSAs predicted by NovumRNA for human cell line UM-UC-3, treated with splicing-perturbation drug 5-Aza-CdR. Samples were taken on day 5,9,13 and 17. Each treatment was performed with two biological replicates, error bars indicate the standard deviation.

**B) Predicted ncTSAs in melanoma cell lines treated with indisulam.** Bar plot shows the mean number of unique ncTSAs predicted by NovumRNA for three human cell lines, 501 MEL, A375, SK MEL 239, treated with splicing-perturbation drugs Indisulam and MS023, DMSO was used as control. Each treatment was performed with three biological replicates, error bars indicate the standard deviation.

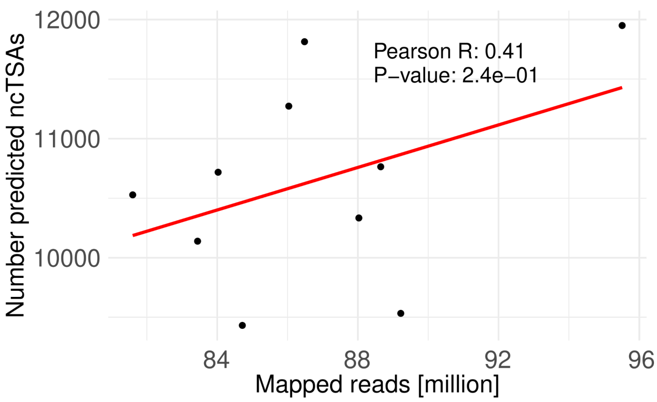


**Figure S3 | Pearson correlation of aligned reads (in millions) and number of predicted ncTSAs with NovumRNA in samples from Ding et al.**

X-axis shows number of aligned reads in millions, aligned to the reference genome (gencode.v41) using hisat2, in UM-UC-3 cell lines. Two Cell lines were treated with methylation inhibitor 5-Aza-2′-deoxycytidine (5-Aza-CdR) and RNA-seq was harvested on day 5,9,13 and 17, including one control. Y-axis shows the number of predicted ncTSAs for each sample.
